# Microvascular homeostasis is compromised in pancreatic islets in a mouse model of beta cell loss and low-grade inflammation

**DOI:** 10.1101/2025.05.05.652204

**Authors:** Luciana Mateus Gonçalves, Isha Shirvaikar, Konstandina Sideris, Elizabeth Pereira, Marjan Slak Rupnik, Joana Almaça

**Author notes:** Correspondence to Joana Almaça. **Conflict-of-interest statement:** The authors have declared that no conflict of interest exists.

## Abstract

Vascular dysfunction is considered a consequence of diabetes. However, in pancreatic islets, some haemodynamic changes occur before the onset of symptoms. The underlying mechanisms driving islet vascular abnormalities have not been fully characterized, but islet pericyte dysfunction appears to be an early event in the pathogenesis of human type 1 diabetes (T1D). It remains to be investigated, however, how abnormal pericyte physiology affects their ability to regulate islet blood flow and vascular permeability. To address this issue, we treated mice with multiple subdiabetogenic doses of the beta cell toxin streptozotocin (STZ; 50mg/kg) and recorded islet vascular responses when animals developed glucose intolerance but were still not diabetic (average fed glycemia <200 mg/dL). At this stage, accompanying increased macrophage density, islet pericytes adopted a myofibroblast-like appearance and interacted closely with endothelial cells expressing high levels of the adhesion molecule ICAM-1. This phenotypical switch of pericytes in STZ-treated mice had functional repercussions: it impacted glucose-induced vasomotor responses *ex vivo* in living pancreas slices and hyperglycemia-stimulated increases in islet blood flow recorded *in vivo* in the exteriorized pancreas. Impaired vasomotor responses were accompanied by enhanced extravazation of a fluorescent dextran (500 kDa) from vessels and accumulation in the interstitial space surrounding islets in STZ-treated mice. Our study indicates that abnormal pericyte function, and compromised capacity to regulate blood flow and vascular integrity, are part of a pathogenic process occurring in islets before diabetes onset, associated with a loss of functional beta cell mass and inflammation.

**Article highlights:**

- Functional and morphological alterations of islet capillaries occur in mice early after multiple low dose STZ treatment;
- Islet pericytes remodel and express higher levels of the myofibroblast marker periostin upon STZ treatment;
- Islet pericyte cytosolic Ca^2+^ and vasomotor responses to high glucose are impaired in living pancreas slices from mice treated with STZ;
- Islet hyperemic responses to increases in glycemia recorded in the exteriorized pancreas are abolished in STZ-treated mice;
- Vascular alterations are associated with a pro-inflammatory environment as islets from STZ-treated mice have more macrophages, an inflamed and leaky endothelium.

## Introduction

Diabetes is a complex immune/metabolic disease during which the body does not produce, secrete or respond to insulin effectively, leading to a predominantly catabolic state of an organism and a rise in blood sugar levels. While T1D is caused by an autoimmune destruction of insulin-producing beta cells in the pancreas(1), type 2 diabetes (T2D) is characterized by insulin resistance and beta cell dysfunction(2). In both forms of diabetes, changes in glucose homeostasis triggers severe complications particularly in the vascular system(3). Although alterations associated with the vasculature are usually considered complications of diabetes, there is accumulating evidence that vascular dysfunction may precede the onset of different endocrine diseases(4), including diabetes. Studies have reported that structural and functional abnormalities of islet blood vessels start during pre-symptomatic stages of the disease(5, 6), but cellular and molecular mechanisms underlying these vascular defects in islets have been poorly characterized.

Islet vascular networks consist of capillary tubes made of endothelial cells covered by pericytes(7, 8). Pericytes act as major regulators of microvascular homeostasis in different tissues(9, 10), but in islets they help regulating endocrine cell function by displaying both trophic and vascular roles(11). Pericytes impact islet beta cell responses to glucose – by enabling acute changes in islet blood flow that occur upon a glucose challenge [functional hyperemia(12, 13)] and by secreting different molecules that potentiate insulin secretion(14, 15). Pericytes have in addition immunoregulatory functions and have recently been proposed to help orchestrating islet inflammation(16, 17). However, it is not known how islet pericytes change in an inflammatory state, and how these changes impact their microvascular functions, such as their ability to regulate local blood flow and vascular permeability.

Previous studies had raised interest in a “microvascular” approach to diabetes(18). It was shown that administering the beta cell toxin STZ to mice as multiple subdiabetogenic doses not only had a direct toxic effect on beta cells, but it also led to an inflammatory reaction against damaged beta cells(19). Because marked hyperglycemia does not develop in the first couple of weeks after low-dose STZ, but animals become glucose intolerant, this animal model mimics a pre-symptomatic (“pre-diabetic”) disease state(20). Islet inflammation develops a few days after STZ, and it is associated with an increase in vascular permeability(21, 22), with alterations occurring mainly at the level of post-capillary venules(18). We used this animal model to examine islet pericyte and capillary responses in an inflammatory setting. The treatment induced striking changes around islet and peri-islet blood vessels – including an increase in perivascular macrophage density, an inflamed endothelium and abnormal pericyte phenotype – which compromised pericyte [Ca^2+^]i and capillary responses, interfering with acute changes in islet blood flow and vascular integrity. Our study thus supports islet vascular dysfunction as an early event in diabetes pathogenesis.

## Methods

### Multiple low-dose STZ mouse model

We administered streptozotocin (STZ; dissolved in sodium citrate buffer (pH 4.5)) to 3-9 months old mice (both genders) as multiple consecutive doses of 50 mg/kg BW for five days by i.p. injections. Control animlas received equivalent volumes of citrate buffer. All experiments were conducted from days 15-23 after treatment start (**Supplementary Figure 1**). To measure [Ca^2+^]i in mural cells, we crossed NG2-Cre mice (*Cspg4*) promoter/enhancer (Jackson labs; #008533) with floxed-GCaMP3 mice (Jackson labs; #029043). Cre negative animals were used in all other experiments. Glucose tolerance tests (2g/Kg BW; i.p., 6h fasting) were performed 10 days after STZ. Experiments were approved by the University of Miami Institutional Animal Care and Use Committee.

Recording mural cell cytosolic Ca^2+^ responses and vasomotion in living pancreas slices Living pancreas slices (150 μm thickness) were prepared from NG2-GCaMP3 mice injected i.v. with a fluorescent lectin (DyLight 649; 70 μL) 10min before sacrifice. Using an upright confocal microscope (Leica SP8), we recorded changes in islet (identified using backscatter) mural cell [Ca^2+^]i and blood vessel diameter induced by glucose (16mM) and epinephrine (10μM). Pericytes were distinguished from SMCs by analyzing their morphology, shape of their cytoplasmic processes (elongated *versus* circular, respectively), location (within islet or covering larger vessels (diameter>10μm) at the islet border or periphery) as previously published(13)(23).

### *In vivo* recording of blood flow and vascular leakage in the exteriorized pancreas

Animals were injected i.v. with 70 μL of fluorescent dextran (500 kDa FITC-Dextran). Animals were anesthetized with isoflurane and a small incision in their abdomen was made to exteriorize their pancreas. The exteriorized pancreas was placed bellow a 3D-printed vacuum window holding a glass coverslip(24) to keep the tissue steady under the confocal microscope. We imaged islet vessels in confocal planes before (baseline) and after glucose (2g or 4g/Kg BW; i.p.) or epinephrine administration (0.9 mg/Kg BW; i.p.), at 15 frames/s for 2 min every 5 minutes. Glycemia was measured before, upon isoflurane and after glucose. Dextran extravazation was determined by quantifying fluorescence intensities in ROIs placed in interstitial regions around the islet periphery, and dividing by fluorescence in islet vessels.

### Automated analysis of capillary blood flow using a Python pipeline

To quantify blood flow, we have adapted a set of interdependent tools to automatize as much as possible the analysis pipeline(25). Detailed analysis is described in Supplementary Experimental Procedures and the complete pipeline is on GitHub: https://github.com/MSlakRupnik/Inst_Speed. In short, we detect “events” which are changes in fluorescent plasma interrupted by dark periods caused by individual red blood cells (RBCs) moving through the vessel (**Supplementary Figure 5**). Each event has been characterized by a start time (t0), its maximal height, and the width at the half of the height (halfwidth, δt), which is our measurement of its duration. The inverse value of the halfwidth of the single event within the ROI of a known size has been used to calculate the mean instant velocity (instantaneous speed; μm/s), normalized to the frame rate of the video. The parameters we used to compare changes in blood flow were the instantaneous speed and the number of events collected for each ROI. We averaged these data for different capillaries for each islet and quantified absolute and relative changes induced by glucose or epinephrine administrations.

### Immunohistochemistry

Two-three weeks after the first injection of STZ or vehicle mice were perfused with 4% PFA and their pancreata collected and cryopreserved for immunohistochemistry. A cohort of mice were injected i.v. with lysine-fixable 40kDa FITC-dextran (10mg/mL). Living pancreas slices were immersed in 4% PFA for 1h and washed with PBS. Detailed information on antibodies used and staining protocols are provided in Supplementary material. Using ImageJ software, the area immunostained for different antigens was quantified and compared to the total area of the region. Mander’s coefficients were used to estimate colocalization between pericyte markers (NG2 and PDGFRβ) and GFP or periostin, in confocal images using the ImageJ plugin “Just Another Co-localization Plugin” (https://imagej.nih.gov/ij/plugins/track/jacop2.html).

### Real-time PCR of sorted islet pericytes

Pancreatic islets were isolated from NG2-GCaMP3 treated with STZ or with vehicle. Islets were processed for quantitative real-time PCR as described(26).

### Statistical analyses

For statistical comparisons we used Prism 10 (GraphPad software) and performed unpaired t-tests when comparing vehicle-treated with STZ-treated mice. One sample t tests were used to determine if average changes in vessel diameter induced by 16G or epinephrine were significantly different from 0. *p*<0.05 was considered statistically significant. Data are presented as individual points, mean ± SEM or in box-and-whiskers plots.

## Results

### Treating mice with multiple low doses of STZ creates glucose intolerance

T1D is a disease of both the endocrine pancreas and of the immune system, and studies have shown that both beta and immune cells are active disease players(27). To examine how the function of islet blood vessels is impacted when beta cells start to fail and an inflammatory setting is present(19), we administered STZ to mice as multiple low doses (50 mg/Kg i.p., 5 days). Non-fasting glycemia started to increase ten days after STZ treatment, although average values remained below 200 mg/dL when experiments were conducted (**Supplementary Figure 1**). By then (two weeks after starting the treatment) animals that received STZ were glucose intolerant and fed plasma insulin levels were lower already one week after STZ (**Supplementary Figure 1)**. Although a significant decrease in beta cell mass occurs in this animal model few days after STZ(28), insulin-positive islets remained two weeks after completion of the treatment (**Figures 1,2**), allowing us to assess changes in the islet microenvironment and vascular function at this stage.

**Figure 1.**
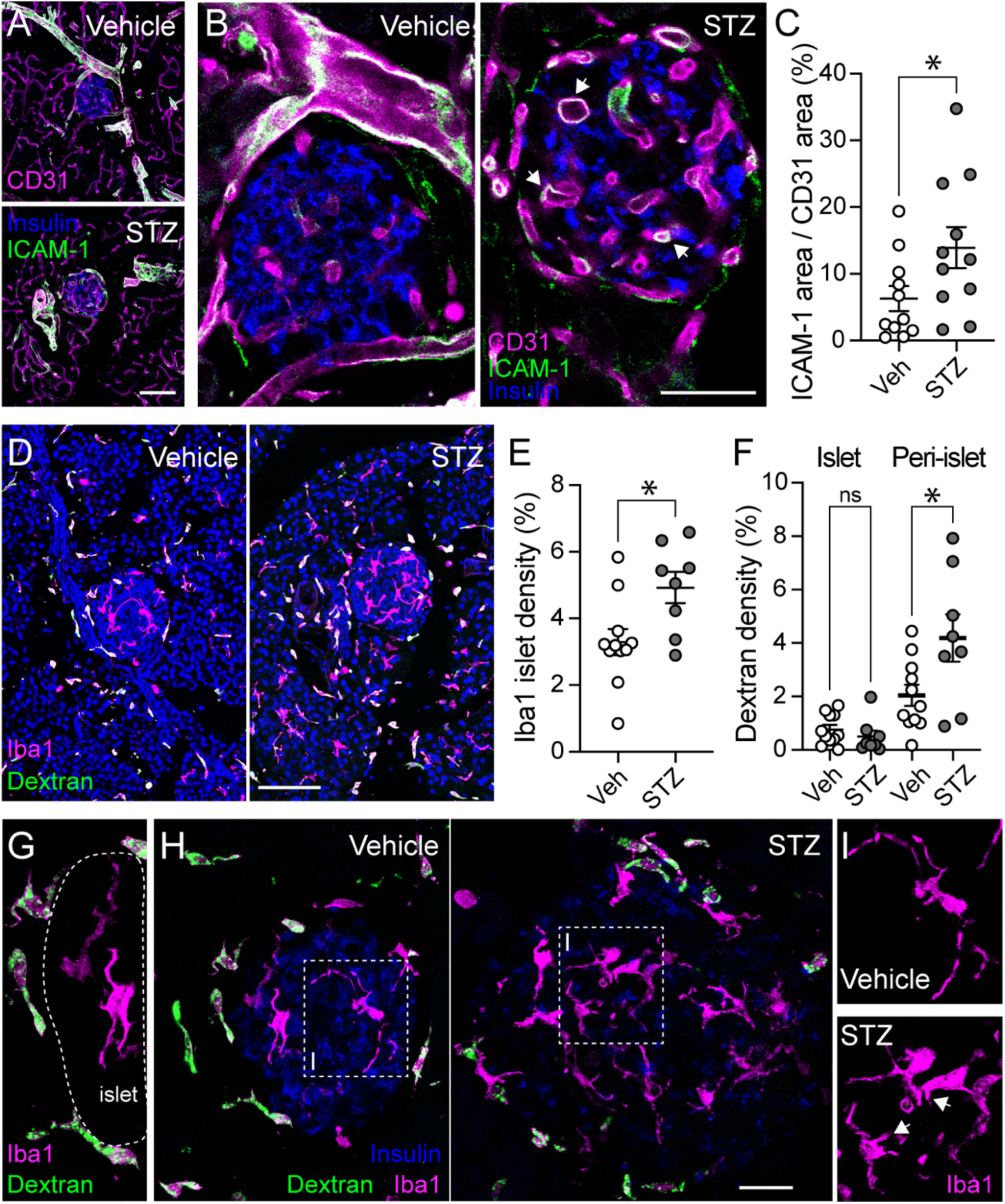
Islets from multiple low-dose STZ treated mice have more macrophages and an inflamed endothelium. (A) Z-projections of confocal images of pancreatic tissue from vehicle- and STZ-treated mice immunostained for CD31 (magenta), ICAM-1 (green) and insulin (blue). Mice were treated with multiple low doses of STZ (50 mg/Kg body weight; 5 days) or with vehicle. (B) Confocal images of islets in vehicle and STZ-treated mice showing insulin (blue), CD31 (magenta) and ICAM-1 (green). (C) Quantification of the area immunostained with ICAM-1 divided by the area immunostained with CD31 in islets from vehicle and STZ-treated mice (n=11 islets/3 mice/group; unpaired t-test; *p*=0.048). (D) Z-projections of confocal images of the pancreas of vehicle- and STZ-treated mice showing macrophages labeled with antibody against ionized calcium-binding adaptor molecule 1 (Iba1; magenta) and dextran (40 kDa FITC-dextan; green). Dextran was injected intravenously in vehicle- and STZ-treated mice 30 min before sacrifice. (E) Quantification of the area immunostained with Iba1 antibody divided by the islet area in vehicle and STZ-treated mice (n=8-11 islets/3 mice/group; unpaired t-test; *p*=0.016). (F) Quantification of the fraction of tissue area labeled with FITC-dextran (inside the islet or within a 20 μm ring that was drawn around each islet (peri-islet)). (G) Dextran is taken up by macrophages at the islet border but not by islet resident ones. (H) Z-projections of confocal images of islets immunostained for Iba1 (magenta), insulin (blue); dextran shown in green. (I) Zoomed images of macrophages within dashed boxes in (H) in islets from vehicle- or STZ-treated mice. Note the change in the shape, process length (arrows) and number of macrophages in STZ islets. Scale bars = 100 μm (A,D), 40 μm (B), 20 μm (H).

**Figure 2.**
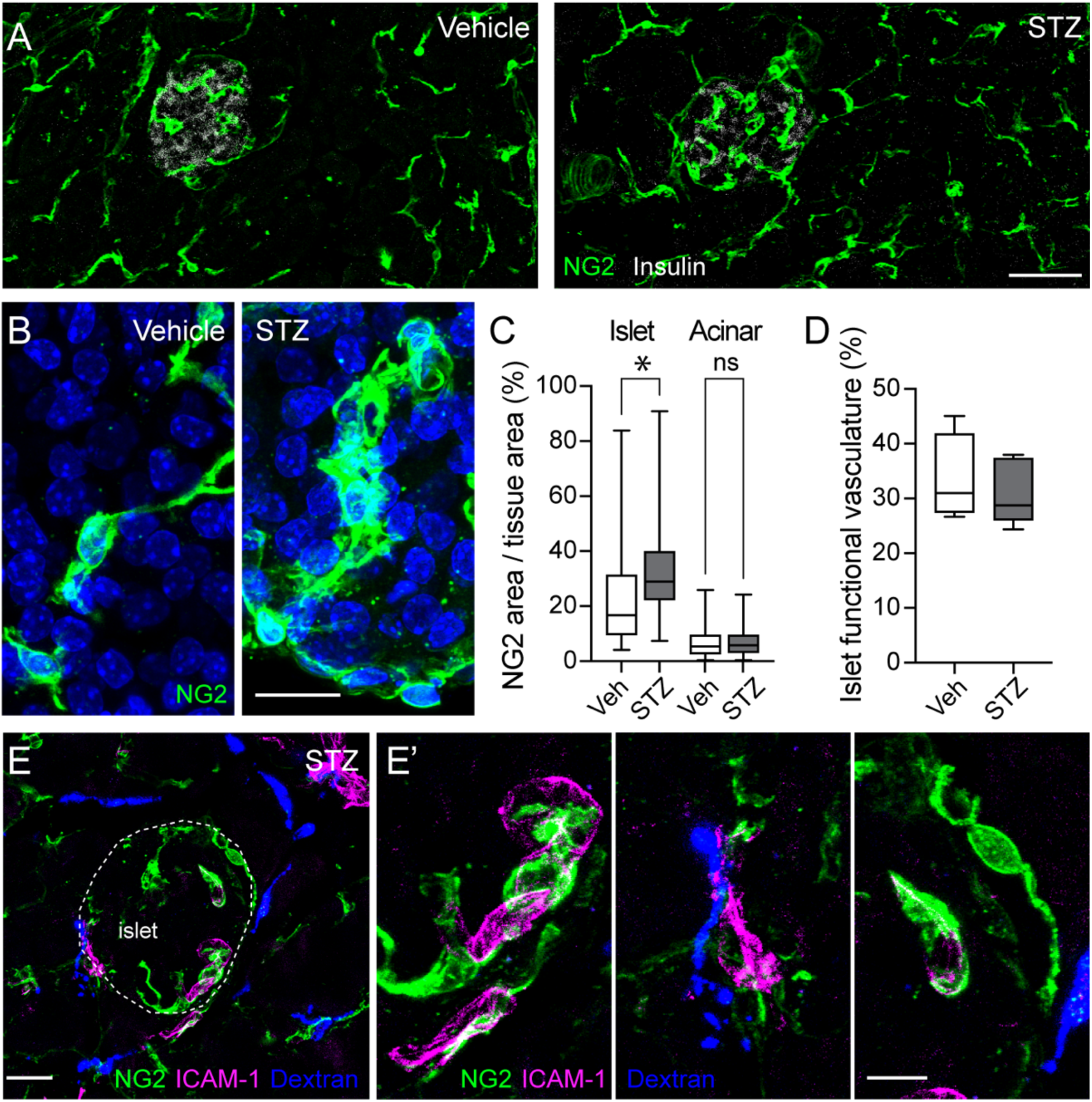
Islet pericytes remodel and acquire an abnormal morphology upon multiple low-dose STZ treatment. (A) Z-projections of confocal images of islets in the pancreas of vehicle- or STZ-treated mice showing beta cells (insulin; gray) and pericytes (labeled with an antibody against the pericyte marker neuron-glial antigen 2; NG2; green). Note the increase in NG2 immunostained area in islets in STZ-treated mice. (B) Zoomed images of pericytes (NG2; green) within islets from vehicle or STZ-treated mice. (C) Quantification of the fraction of tissue area (islet or acinar tissue) immunostained with NG2 (n=25-32 islets/4 mice/group; one-way ANOVA followed by Tukey’s multiple comparisons test; \**p*=0.02); (D) Quantification of islet functional vasculature, i.e. fraction of the islet area labeled with a fluorescent lectin from *Lycopersicon esculentum* (DyLight649), which was injected into the tail vein of STZ or vehicle treated animals (n=11-13 islets/4 mice/group; unpaired t-test; *p*=0.35). (E) Z-projection of confocal images of an islet in the pancreas of a STZ-treated mouse immunostained for NG2 (green), ICAM-1 (magenta) and dextran-labeled macrophages (blue). (E’) Zoomed images of regions in (E) showing dextran-positive macrophages and pericytes with hypertrophied processes interacting with ICAM-1-labeled endothelial cells. Scale bars = 50 μm (A), 10 μm (B,E’), 20 μm (E).

### Islets from STZ-treated mice have more macrophages and an inflamed endothelium

Studies had shown that treating mice with multiple low doses of STZ caused an inflammatory process in islets that started a few days after completion of the treatment(19). To examine the magnitude of islet inflammation, we first assessed the expression of intercellular adhesion molecule 1 (ICAM-1; known as CD54) in islets from animals treated with STZ or with vehicle. ICAM-1 is an adhesion molecule crucial for immune cell recruitment into tissues(29), and one of the genes significantly upregulated in islets from mice treated with different regimens of STZ(30). While ICAM-1 is expressed by CD31-positive endothelial cells lining some large vessels in the exocrine tissue and at the islet periphery (**Figure 1A**), capillaries inside islets from vehicle-treated mice were devoid of ICAM-1 (**Figure 1B**). In contrast, several islet endothelial cells in STZ-treated mice stained positive for ICAM-1 (**Figure 1B**, arrows), and there was a significant increase in ICAM-1 vascular levels in islets from STZ-treated animals (**Figure 1C**), indicative of an inflamed endothelium at this stage.

We then examined whether treatment affected the density and phagocytic capacity of macrophages, the predominant immune cell type in pancreatic islets at steady state(31). Macrophages are the first immune cell type to infiltrate islets in mice treated with multiple doses of STZ(32, 33). Macrophages in the endocrine and exocrine pancreas were visualized with an antibody against Iba1 (**Figure 1D**). In line with previous studies, we found a significant increase in the density of Iba1-labeled macrophages in islets from STZ-treated mice(**Figures 1D,E,H**). Besides an increase in number, we also noticed that the shape of islet resident macrophages was different in STZ-treated mice, and these cells exhibited either fewer or shorter cytoplasmic projections (**Figure 1I**). In addition, we injected a 40kDa fluorescent dextran in the tail vein and let it circulate for 30 min before sacrificing the animals. This dextran is taken up by macrophages in the exocrine tissue of the pancreas but not by islet resident ones (**Figure 1D**).

Indeed, dextran-labeled macrophages could be seen in the periphery of islets from vehicle- and STZ-treated mice but they did not penetrate the islet parenchyma (**Figures 1G,H**). These data are in line with previous studies showing that islet macrophages are phenotypically different than those at the islet border or in acinar tissue(34). The density of dextran-labeled cells was significantly higher at the border of islets in STZ-treated mice (**Figure 1F**). In summary, upon STZ, the islet endothelium is inflamed and there are more macrophages inside and at the periphery, indicative of an ongoing inflammatory process.

### Islet pericytes remodel and acquire an abnormal morphology upon STZ treatment

A recent study showed that pericytes participate in the regulation of islet inflammation by producing and sensing different inflammatory cytokines(17). Remodeling of islet pericytes and abnormal ultrastructure had been reported in animals that developed hyperglycemia upon STZ(7, 35), but it is not known what happens to this cell population when marked dysglycemia has not yet developed. We first asked whether there were changes in the density of pericytes in endocrine and exocrine pancreas of mice treated with multiple doses of STZ. Pericytes were labeled with an antibody against the pericyte marker neuron-glial antigen 2 (NG2). We noticed that there was a significant increase in the area immunostained with NG2 antibody in islets from STZ-treated mice but not in the surrounding exocrine tissue (**Figures 2A,C**). This increase in the density of islet pericytes was not due to an increase in the number of cells expressing NG2, as this parameter was not different between STZ- or vehicle treated animals (**Supplementary Figure 2**). To determine whether changes in islet pericyte density reflected alterations in the density of vascular structures in islets, we injected a fluorescent lectin into the tail vein of STZ or vehicle treated animals. We found that the density of functional blood vessels in islets was not different between the groups (**Figures 2D,4A**), in contrast with previous data showing that islet capillaries were less numerous (25% less) and appeared collapsed in islets from mice treated for 5 days with STZ(36).

Notably, pericytes in islets from STZ-treated animals had an abnormal shape, and some of them had lost the typical elongated morphologies that characterize this cell population (**Figure 2B**). In some regions, islet pericytes had atypical spreading and short protrusions of their cytoplasmic processes. Interestingly, these pericytes with hypertrophied processes and abnormal morphology were closely associated with ICAM-1 expressing endothelial cells (**Figure 2E**), further supporting pericytes’ potential involvement in the ongoing inflammatory process(16, 17). In summary, islet pericytes remodel and acquire an abnormal morphology upon multiple low-dose STZ treatment.

### Pericytes adopt a myofibroblast-like phenotype in islets from STZ-treated mice

Because inflammation is usually linked to fibrosis(37) and pericytes can turn into myofibroblasts(26), we then assessed whether a similar shift in the phenotype of islet pericytes occurred upon STZ. To label myofibroblasts (also known as activated stellate cells), we used an antibody against periostin, a secreted matricellular protein involved in cellular adhesion and organization of collagen in the heart(38) and islets(26). In vehicle-treated, mice periostin could be detected at the islet border but rarely inside the endocrine parenchyma (**Figure 3A**). However, in STZ-treated animals the density of periostin at the islet-exocrine interface was much higher, and periostin-positive structures could be detected inside the islet (**Figure 3B**). Importantly, while periostin was not expressed by intra-islet pericytes in vehicle-treated mice (although some pericytes at the islet border expressed periostin; **Figure 3C**), the degree of colocalization of NG2 and periostin was significantly higher in STZ-treated animals (**Figures 3D,E**). These data indicate that in STZ-treated mice a subset of islet pericytes (∼40%) has adopted a myofibroblast-like phenotype.

**Figure 3.**
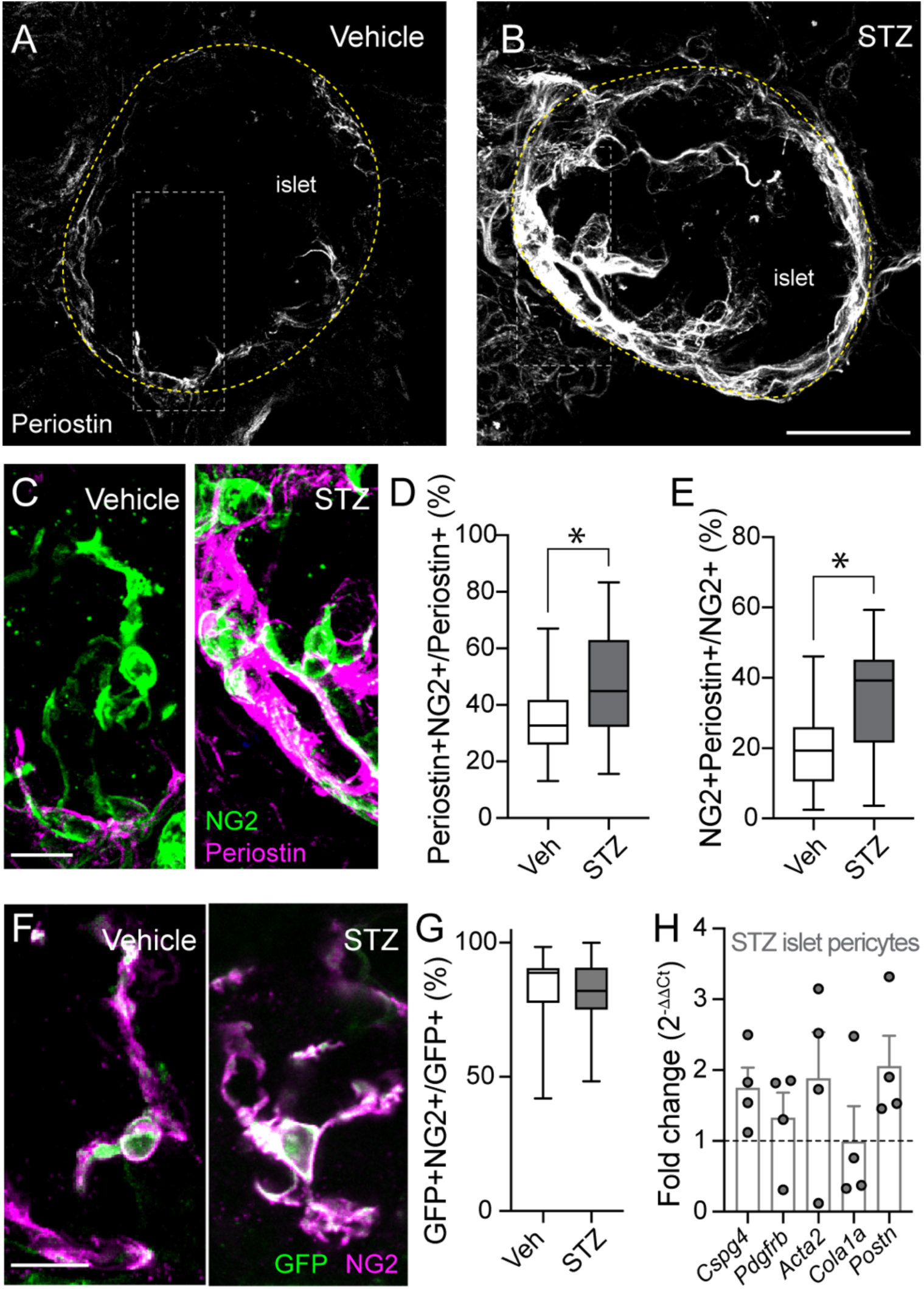
Pericytes adopt a myofibroblast-like phenotype in islets from STZ-treated mice. (A,B) Z-projections of confocal images of islets in the pancreas of vehicle- (A) or STZ-treated mice (B) showing periostin immunostaining (gray). Note increased periostin immunostaining around the islet upon STZ treatment. (C) Zoomed images of regions within dashed white rectangles in (A,B) showing NG2-labeled pericytes (green) and periostin (magenta). (D,E) Mander’s coefficients estimating colocalization between NG2 and periostin in confocal images of islets treated with vehicle or STZ (n=22-27 islets/3-4 mice/group; unpaired t-tests; *p*≤0.01). (F) Z-projections of confocal images of pericytes in islets from NG2-GCaMP3 mice treated with vehicle or STZ showing GFP (green) and NG2 (magenta) immunostaining. Note loss of elongated shape of pericytes in STZ islet. (G) Mander coefficient estimating colocalization between GFP and NG2 (n=19-28 islets/4-6 mice/group; unpaired t-test; *p*=0.996). (H) Fold change in mRNA levels in sorted islet pericytes (GFP-expressing cells) from STZ-treated mice compared to levels of the same genes in pericytes from vehicle-treated animals (n=4 mice/group). Genes examined include different pericyte and (myo)fibroblast markers (*Cspg4*, *Pdgfrb*, *Acta2*, *Cola1a* and *Postn*). Scale bars = 40 μm (A,B), 10 μm (C,F).

**Figure 4.**
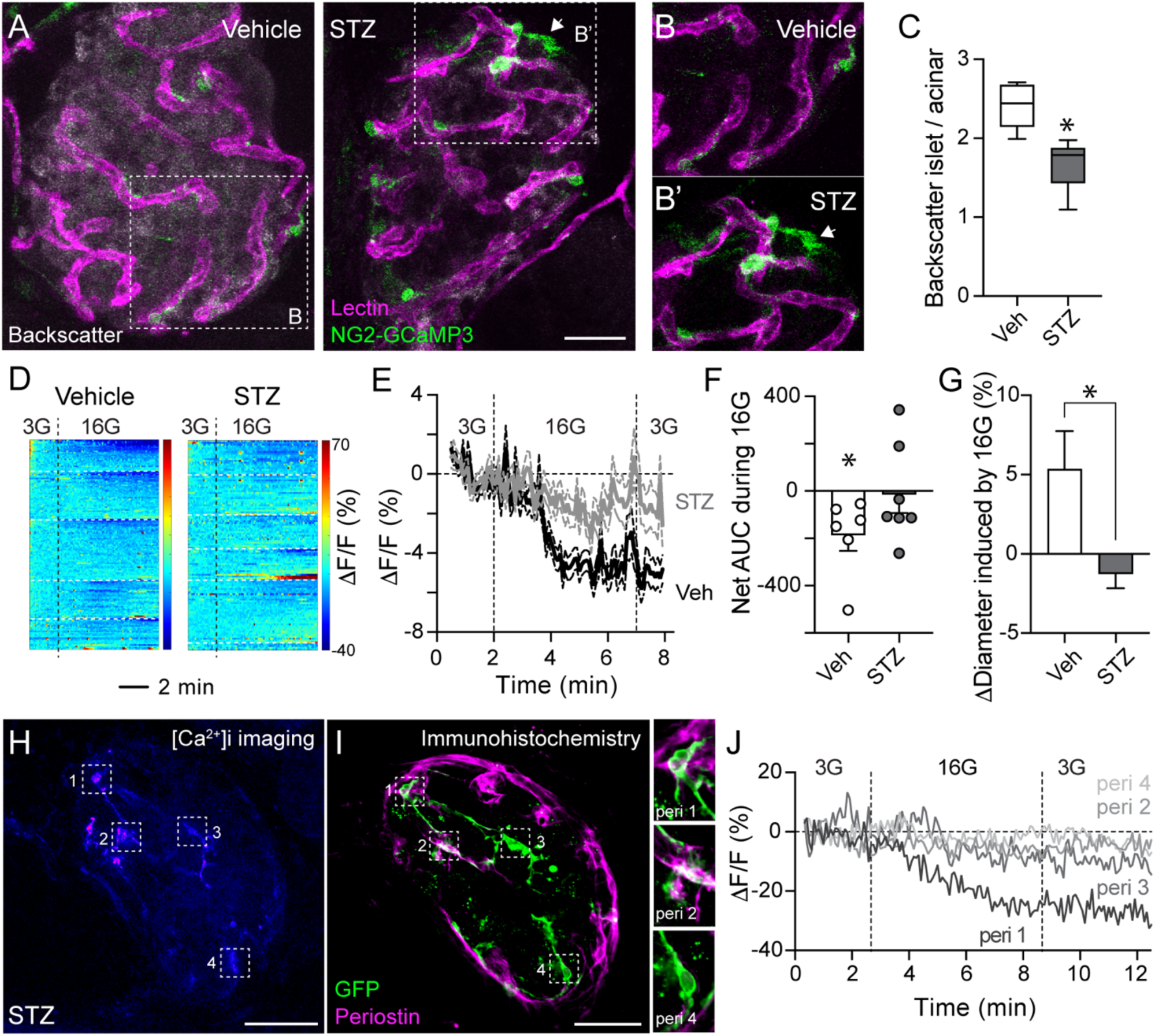
Islet vasomotor responses to glucose are impaired *ex vivo* in living pancreas slices from STZ-treated mice. (A) Z-projections of confocal images of islets in living pancreas slices produced from mice expressing GCaMP3 (green) under the *Cspg4* promoter (NG2-GCaMP3 mice), treated with either vehicle or STZ. Animals were injected with fluorescent lectin (DyLight649; magenta) 10 min before sacrifice and slice production. Islets in slices can be visualized because of the backscatter signal that dense-core granules in endocrine cells produce (backscatter; gray). (B,B’) Zoomed images of regions in islets in (A) showing GCaMP3-expressing islet pericytes (green) and islet capillaries (magenta). Arrow indicates a pericyte in a STZ islet detached from a capillary. (C) Quantification of the ratio of backscatter signal in islets divided by signal produced by surrounding acinar tissue. STZ treatment leads to a decrease in backscatter signal (n=11-13 islets/ 4-5 mice/ group; unpaired t-test; *p*=0.0002). (D) Heatmaps showing changes in pericyte [Ca^2+^]i (GCaMP3 fluorescence; 1F/F (%)) induced by increasing glucose concentration from 3 mM to 16 mM (16G; dashed vertical line indicates when 16G was applied). Dashed horizontal white lines separate pericyte responses from different mice in each group (n=5 vehicle-treated; 6 STZ-treated). (E) Traces (average (thicker line) ± SEM (thinner lines)) showing relative changes in GCaMP3 fluorescence levels in islet pericytes induced by 16G in islets from mice treated with vehicle (black) or STZ (gray). (F) Quantification of the net area under the curve of fluorescence traces as in (E) showing the effect of 16G on islet pericyte [Ca^2+^]i. Each dot represents the average of 4-46 pericytes per islet from 6 vehicle- and 7 STZ-treated mice (one sample t-test compared to theoretical mean of 0; *p*=0.04). (G) Quantification of the relative change in islet capillary diameter induced by high glucose (% initial diameter; n=46 capillaries/4 islets/3 mice; *p*=0.01 unpaired t-test). (H,I) Islet pericyte [Ca^2+^]i responses to 16G were recorded (H; pseudo color scale) and the same slice was then fixed and immunostained (I) for GFP (green) and periostin (magenta) to correlate pericyte calcium responses to glucose with their phenotype. Zoomed images of pericytes withing dashed squares in (H,I) are shown on the right in (I). (J) Traces of relative changes in GCaMP3 fluorescence induced by 16G of islet pericytes shown within dashed squares in images in (H,I). Scale bars = 20 μm (A), 40 μm (H,I).

To further examine phenotypical and functional changes associated with islet pericytes, we generated mice that express a fluorescent reporter (and calcium indicator; GCaMP3) in mural cells [pericytes and smooth muscle cells; using Cre recombinase under the NG2 (*Cspg4*) promoter;(12)]. Treating these animals with STZ did not affect the proportion of GCaMP3 (GFP) that is expressed by pericytes (and colocalizes with NG2 or PDGFRβ; ∼80% in vehicle and STZ-treated islets for colocalization estimates between GFP/NG2 and GFP/PDGFRβ; **Figures 3G,F, Supplementary Figure 2**). We isolated islets from vehicle and STZ-treated mice and sorted GFP-expressing islet pericytes by flow cytometry (**Supplementary Figure 2**). We then examined the levels of various genes encoding pericyte markers (*Cspg4, Pdgfrb, Acta2*), ECM and myofibroblast markers (*Cola1a, Postn*) by real-time qPCR. We found that transcript levels of genes encoding NG2 (*Cspg4*) and periostin (*Postn*) were higher in pericytes sorted from islets isolated from all STZ-mice (n=4) when compared to levels in vehicle-treated animals (**Figure 3H**), further supporting a phenotypical switch of islet pericytes.

### Islet vasomotor responses to glucose are impaired in living slices from STZ-treated mice

To examine the potential impact of the phenotypical shift on pericyte activity and vasomotion in islets, we produced living pancreas slices from NG2-GCaMP3 two weeks after STZ treatment. Not surprisingly, STZ treatment resulted in a significant decrease in islet backscatter signal (**Figures 4A,C**), which is an intrinsic indicator of the amount of mature insulin granules within islet endocrine cells(39). As shown in Figure 2D, STZ treatment did not affect the density of functional blood vessels in islets or the average basal capillary diameter (which remained around 5 μm) but islet capillary diameter was more heterogenous in STZ-treated mice (**Figures 4A,B**, **Supplementary Figure 3**). Some pericytes in STZ islets appeared detached from capillaries (arrow in **Figures 4A,B**).

We had previously shown that increasing extracellular glucose concentration led to a decrease in [Ca^2+^]i in a subset of pericytes in mouse and human islets and an increase in islet capillary diameter (13, 40). We then asked whether islet vascular responses to glucose were preserved in our STZ model. To this end, we recorded islet pericyte [Ca^2+^]i and capillary responses to a switch in extracellular glucose concentration from 3 mM to 16 mM (16G) in pancreas slices from vehicle- or STZ-treated mice. We found that increasing glucose concentration decreased [Ca^2+^]i in a subset of pericytes in islets from vehicle-treated but not as much in islets from STZ-treated mice (**Figures 4D-F**). When we averaged pericyte [Ca^2+^]i responses from different islets and mice, high glucose led to a ∼5% decrease in GCaMP3 fluorescence levels in pericytes in vehicle-treated but only to ∼1% decrease in STZ-treated animals (**Figure 4E**). This decrease in islet pericyte [Ca^2+^]i upon stimulation with high glucose was accompanied by a ∼5% average increase in islet capillary diameter in slices from vehicle-treated mice, but capillaries from islets of STZ-treated mice did not dilate (**Figure 4G**). These data indicate that vascular responses to high glucose are impaired in STZ islets.

We then examined whether impaired pericytic responses to glucose in STZ islets depended on their switch of phenotype and periostin expression. To this end, we recorded pericyte [Ca^2+^]i responses to high glucose ([Ca^2+^]i imaging; **Figure 4H**) and then performed immunohistochemistry on the same slice for GFP and periostin (**Figure 4I**). Recorded pericytes can be nicely matched with subsequent immunostaining images (**Supplementary Figure 4**).

We found that pericytes in islets from STZ-treated mice that did not express periostin (e.g. pericytes #3,4; **Figures 4H,I**) still exhibited abnormal responses to high glucose (**Figure 4J**). These data indicate that islet pericytes were overall dysfunctional in STZ-treated mice, independently of their periostin expression and myofibroblast phenotype.

### Arterioles constrict upon epinephrine, but islet capillaries are stiff after STZ treatment

To assess whether STZ treatment affected only islet vascular responses to glucose or instead had a more general effect and compromised responses to other vasoactive stimuli, we stimulated slices with the sympathetic agonist epinephrine (Epi; 10 μM). We had previously reported that the microvasculature in mouse and human islets was very responsive to different sympathetic agonists such as epinephrine, norepinephrine and phenylephrine(12, 13, 23, 40). In agreement with our previous studies, in vehicle-treated animals, epinephrine administration led to an increase in [Ca^2+^]i in islet pericytes and smooth muscle cells (SMCs) around arterioles present in the vicinity of islets (**Figure 5A**). Epinephrine activated islet pericytes and triggered a ∼15% increase in average GCaMP3 fluorescence levels in these cells (**Figure 5B**) but had an even more powerful effect on SMCs calcium (∼60% average increase in GCaMP3 fluorescence; **Figure 5C**). In vehicle-treated mice, changes in [Ca^2+^]i in islet pericytes and in SMCs were accompanied by changes in vessel diameter, and epinephrine led to a ∼5% decrease in islet capillary diameter and to an even more pronounced constriction of islet arterioles (∼20% decrease in vessel diameter; **Figure 5D**).

**Figure 5.**
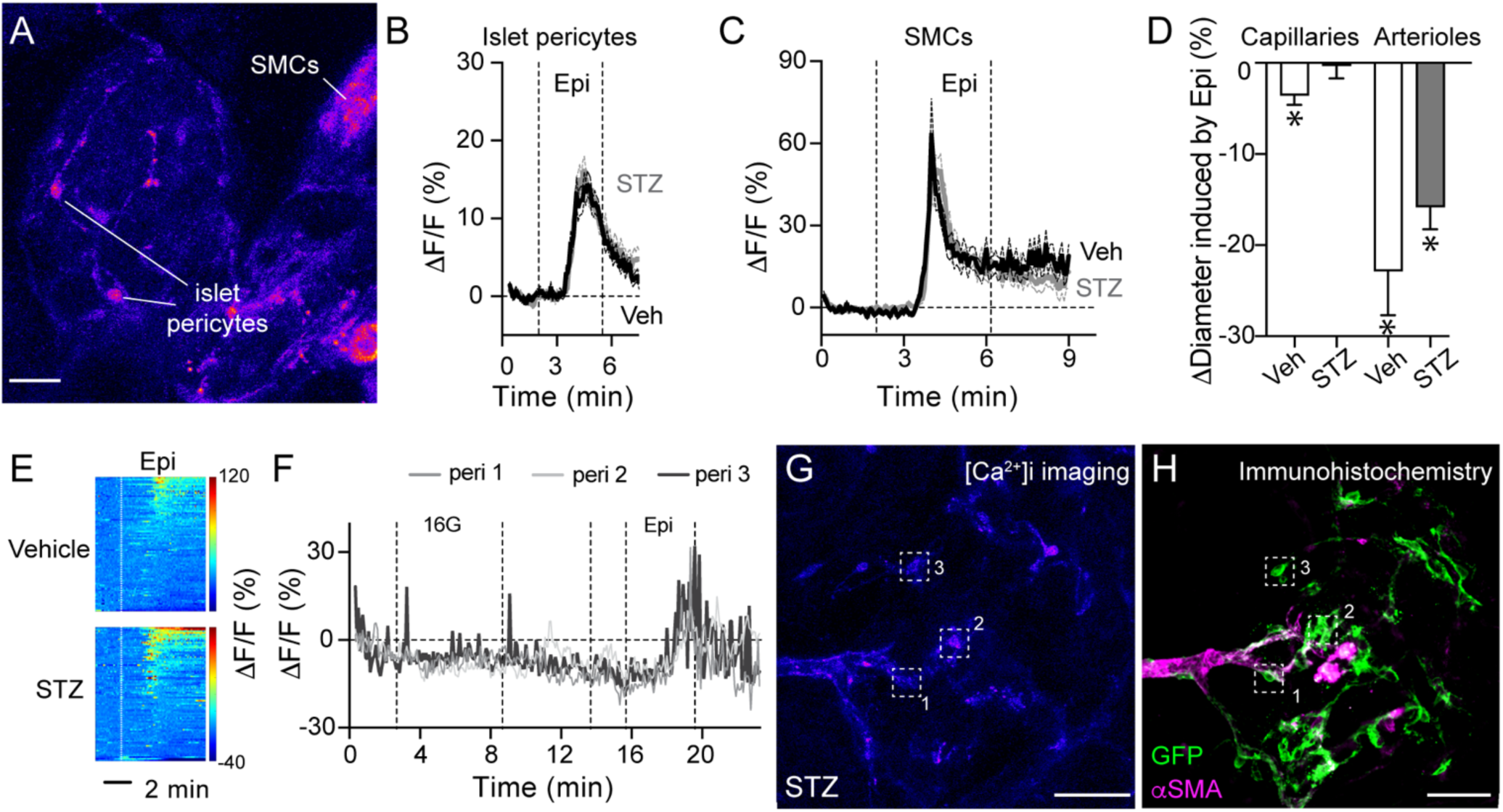
Arterioles constrict upon epinephrine, but islet capillaries are stiff after STZ. (A) Z-projection of confocal images of GCaMP3 expressing mural cells (pericytes and smooth muscle cells (SMCs)) in islets from NG2-GCaMP3 mice. Image is shown in a pseudo color scale. (B,C) Traces (average (thicker line) ± SEM (dashed thinner lines)) showing relative changes in GCaMP3 fluorescence induced by epinephrine (epi; 10 μM) in islet pericytes (B; n=224-235 pericytes/15islets/6-7mice) and SMCs (C; n=18-53 SMCs/3-6islets/2-5mice) from mice treated with vehicle (black) or STZ (gray). (D) Quantification of relative changes in islet capillary and arteriole diameter induced by epinephrine (% initial diameter; n=60-68 capillaries and 4-7 arterioles/6 islets/4 mice; one sample t-test compared to theoretical mean of 0; *p*=0.001). (E) Heatmaps showing changes in pericyte [Ca^2+^]i (GCaMP3 fluorescence; ΔF/F (%)) induced by epinephrine (dashed vertical lines indicate when epi was applied). Each line in the heatmap is a different pericyte (from 6-7 vehicle- or STZ-treated mice). (F-H) Traces of relative changes in GCaMP3 fluorescence induced by 16G and Epi (F) of islet pericytes shown within dashed squares in images in (G,H), during calcium imaging (G) and after immunostaining (H) for GFP and αSMA (magenta). Scale bars = 20 μm (A) 40 μm (G,H).

We then examined mural cell and capillary responses to epinephrine in islets from animals treated with STZ. We found that pericyte and SMCs [Ca^2+^]i responses to epinephrine were preserved in islets from STZ-treated animals (**Figures 5B,C,E**). Indeed, even those islet pericytes that did not decrease their [Ca^2+^]i levels upon high glucose stimulation (**Figure 5F**) still displayed a potent response to epinephrine (**Figures 5F-H**). Interestingly, although pericyte responses appear normal, capillaries in STZ-treated islets did not constrict upon stimulation with this adrenergic agonist (**Figure 5D**). In contrast to islet capillaries, arterioles in the islet periphery were still capable of constricting upon epinephrine, which induced a significant reduction in their diameter (**Figure 5D**). Our data indicate that, unlike in vehicle-treated mice, there is a dissociation between pericyte [Ca^2+^]i and changes in capillary diameter in STZ-treated islets: while islet pericyte [Ca^2+^]i responses to epinephrine were preserved, islet capillaries became stiff with impaired vasomotor capacity in STZ-treated animals. The changes observed were specific to the intra-islet microvasculature as arterioles in the islet periphery remained responsive.

### Hyperglycemia and epinephrine induce acute changes in blood flow in islets in the exteriorized mouse pancreas

Studies have shown that islet blood flow is disturbed during conditions of impaired glucose tolerance and diabetes(41). We hypothesized that remodeling of the islet vasculature and impaired pericyte and capillary responses upon STZ treatment *ex vivo* affected islet blood flow and perfusion *in vivo*. To record islet blood flow in control mice, we injected a fluorescent dextran (150 kDa FITC-dextran) into the mouse tail vein before exteriorizing and immobilizing the pancreas with a 3D-printed vacuum window(24) (**Figure 6A**). Dextran injection allowed us to visualize vascular trees in islets (which could be detected using backscatter; **Figure 6B**). We then recorded blood flow in islets under basal conditions for 30 min, injected glucose i.p. (2 g/kg BW) and followed changes in flow for 45 min, followed by an injection of epinephrine i.p. (0.9 mg/kg BW) and recorded for an additional 10 min period.

**Figure 6.**
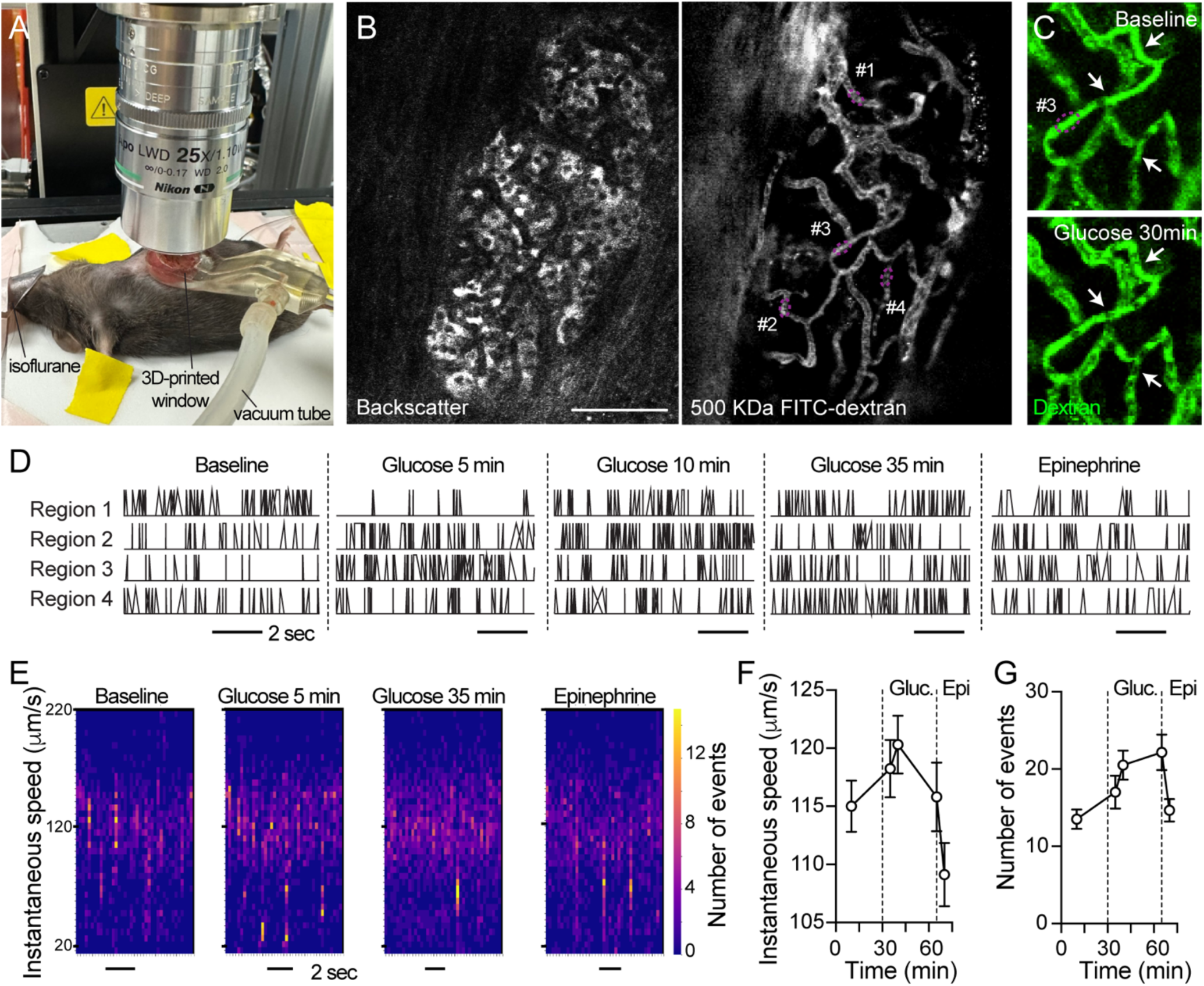
Glucose increases while epinephrine decreases blood flow in islets in the exteriorized mouse pancreas. (A) Exteriorization of the pancreas of an anesthetized mouse for *in vivo* imaging of blood flow. Gentle suction by connecting to a vacuum pump allows for flush contact of tissue with glass coverslip on a 3D-printed window and the best focus for recording. (B) Islets can be found in the pancreas using the backscatter signal (left panel). Blood vessels were labeled with an intravenous injection of 150 kDa FITC-dextran (right panel). Blood flows at different speeds through islet capillaries under basal conditions. (C) Zoomed images of capillaries in region #3 of the islet shown in (B) before (baseline) and 30 min after i.p. injection of 20% glucose (2 g/kg BW i.p.; at this moment, glycemia was 521 mg/dL). Note increase in density of red blood cells (RBCs) – shadows that appear within vessel lumens – in several capillaries (arrows). (D) Movement of RBCs through different capillaries in islet shown in (B) were recorded under basal conditions, at different time points after i.p. injection of glucose (5, 10 and 35 min after glucose), followed by i.p. injection of epinephrine. RBC movement over the course of the videos were converted to binary events and graphed to show the event density and frequency across time. At each time point, 0 was given for no events (no blood flow) and 1 was given when events occurred (blood flowing). Scale bar is equivalent to 2 seconds. (E) Instantaneous RBC speed was calculated using the Python pipeline described in Methods section. Representative heatmaps showing instantaneous speed estimates on the y-axis and the x-axis denotes the time of the recording. At each timepoint, the number of events at each instantaneous speed value is depicted with the color gradient displayed on the right side. Bright yellow is a higher number of events and dark blue is the lowest number of events. The baseline, 5 minutes post glucose, 35 minutes post glucose, and 5 min post epinephrine recordings are compared in this figure. (F,G) Mean ± SEM of instantaneous speed (F) or total number of events (G) through all capillaries in different regions in islet shown in (B) at different time points after glucose and epinephrine. Dashed lines indicate administration of stimuli. Scale bar = 50 μm (B).

To quantify blood flow, we adopted a Python-based analysis pipeline which measures changes in fluorescence that occur in the lumen of vessels when red blood cells (RBCs, which are dark spots in **Figure 6C; Supplementary Movie 1**) cross and interrupt the fluorescence signal in regions of interest (ROIs) positioned on those vessels (25). Briefly, these changes in fluorescence can be converted into measures of instantaneous speed and number of events in ROIs placed on different islet capillaries (see “Experimental procedures” for details and **Supplementary Figure 5**). We noticed that blood flow through different islet capillaries was heterogenous under basal conditions (**Figure 6B; Supplementary Movie 1**), with some capillaries already showing high RBC event density in baseline videos (e.g. regions 1 and 4; **Figure 6D**). Some capillaries that had initially low basal blood flow exhibited a significant increase in event frequency upon glucose stimulation (e.g. region #3; **Figures 6C,D; Supplementary Movies 1,2**). Glucose administration led to an overall increase in islet blood flow, which was characterized by both a rise in average instantaneous speed during the first 15 min (**Figures 6E,F**) and on the number of events for the full period of 45 min (**Figures 6E,G**). These data are in line with previous studies showing that glucose increases blood flow in islets in the pancreas(42, 43) and when they are transplanted into the anterior chamber of the mouse eye(13), a physiological response known as functional hyperemia. Epinephrine administration, in turn, reduced blood flow in islets in the pancreas (**Figures 6D-G**). These data indicate that our *in vivo* imaging platform and analysis pipeline allow us to readily detect acute changes in islet blood flow induced by phyiological stimuli, such as glucose and epinephrine.

### Hyperemic responses to glucose are impaired in islets from STZ-treated mice *in vivo*

Because pericytes are major regulators of functional hyperemic responses in islets induced by glucose(13) and because vasomotor responses to glucose are impaired in islets from STZ-treated mice *ex vivo* (**Figure 4**), we then asked whether this glucose-dependent increase in islet blood flow would still occur upon STZ treatment . We noticed that STZ treatment slightly increased basal islet blood flow, characterized by more events at different speeds (**Figures 7A,C**). In animals treated with vehicle, glucose led to a significant increase in instantaneous speed of RBCs in islet capillaries during the initial 15min after stimulation (**Figures 7B,D,E**) but not in capillaries in the acinar tissue (**Supplementary Figure 5**). This glucose-induced acute increase in instantaneous RBC speed in islets was abolished in mice that received STZ, and glucose led instead to a decrease in blood flow in both islets and acinar tissue in these animals (**Figures 7B-E,Supplementary Figure 5**). In contrast, epinephrine-induced decrease in blood flow was still observed in islets from STZ-treated mice, most likely due to preserved SMCs [Ca^2+^]i responses and arteriole constriction in islets from these mice (**Figure 5**).

**Figure 7.**
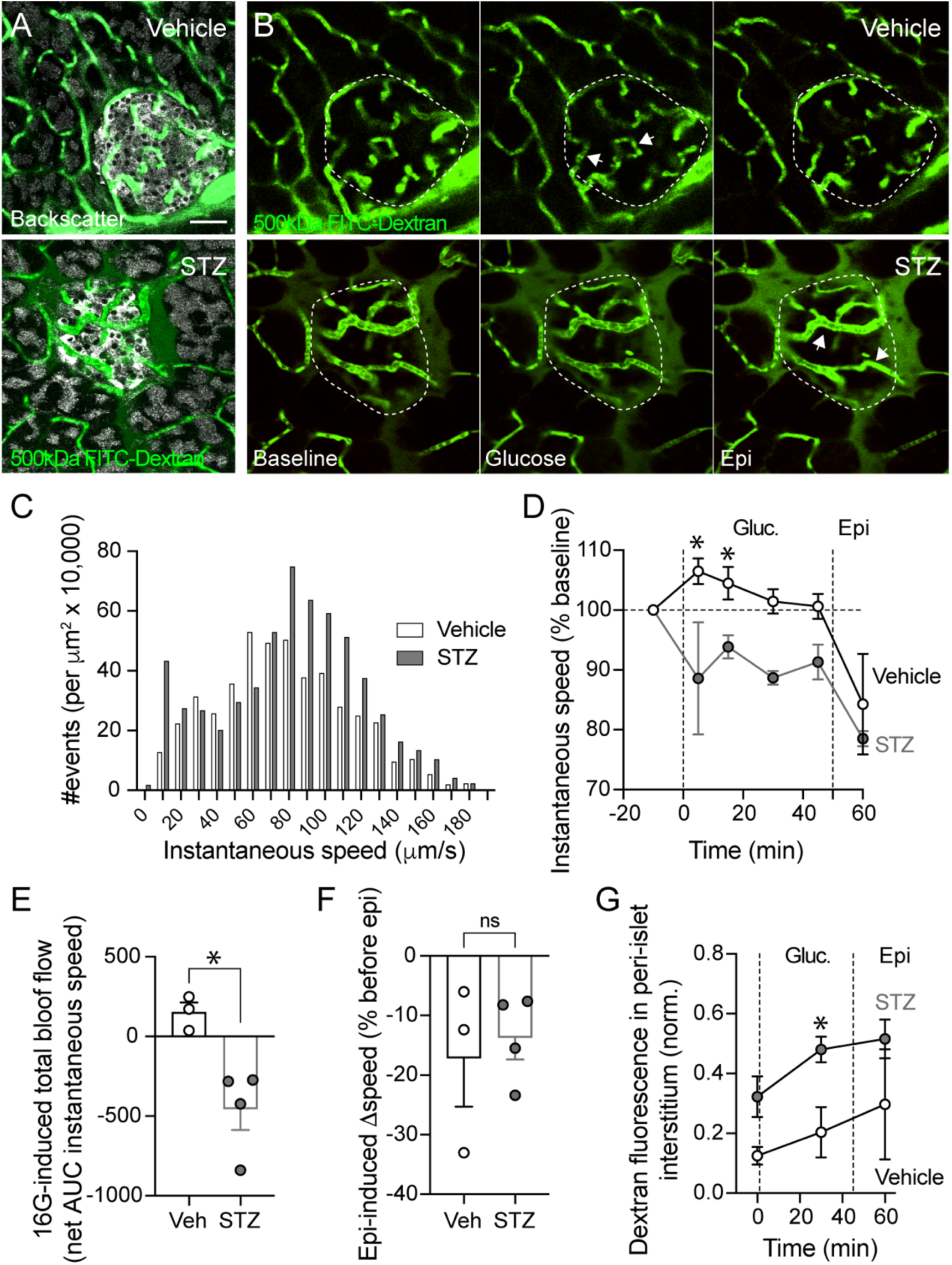
Hyperemic responses to glucose are impaired in islets from STZ-treated mice. (A) Confocal images of islets in the exteriorized pancreas of a vehicle- or a STZ-treated mouse. Blood vessels are labeled with FITC-dextran (green; 500kDa) and islets are identified with backscatter (gray). Scale bar = 50 μm. (B) Snapshots of islets shown in (A) showing dextran labeling before (baseline), 30 min after injecting intraperitoneally 40% glucose (4 g/kg glucose i.p.; glycemia reached 600 mg/dL for both groups), and 10 minutes post epinephrine. Arrows indicate vessels where blood flow changed. (C) Histogram showing the number of events (normalized to the vessel area and multiplied by an average islet area of 10,000 μm^2^) with instantaneous speeds ranging from 0-180 μm/sec. (D) Quantification of relative changes in instantaneous speed (shown as a % of baseline speed) of blood flowing through capillaries in islets in mice treated with STZ (n=4 mice; gray) or vehicle (Veh; n=3 mice; black). Values were measured at 5, 15, 30, and 45 minutes after i.p. glucose, and 10 minutes post epinephrine (n=3-4 mice; multiple unpaired t test; \**p*< 0.05). (E) Quantification of the net AUC of traces shown in (D) reflecting changes in instantaneous speed in islet capillaries induced by glucose (unpaired t-test; *p*<0.05). (F) Quantification of relative changes in instantaneous speed induced by epinephrine in islet capillaries from vehicle- and STZ-treated mice (unpaired t-test; *p*>0.05). (G) Quantification of dextran extravazation to the interstitial space surrounding islets (fluorescence levels normalized to those in islet vessels; n=3-4 islets/group (1 islet/mouse); multiple unpaired t test; \**p*< 0.05).

Interestingly, we noticed that the FITC-dextran (500kDa) we had used to label the vasculature in these experiments extravazated from the vessels towards the interstitial space surrounding islets in the pancreas of STZ-treated mice (**Figures 7A,B**). Dextran leakage progressively increased with recording time and its intensity in the islet periphery was significantly higher in animals treated with STZ (**Figure 7G**). Leakage was not limited to regions surrounding islets, as some dextran extravazation could also be observed from exocrine capillaries in animals that had received STZ (**Supplementary Movies 3,4**). In summary, our study shows that STZ treatment impacts the function of islet pericytes, acute regulation of islet blood flow and vascular permeability.

## Discussion

In this manuscript we examined the function of the islet microvasculature early after administering the beta cell toxin STZ in multiple sub-diabetogenic doses. This regimen of STZ administration leads to a progressive decline in functional beta cell mass concomitantly with the development of an early inflammatory process in islets. Importantly, in the first two weeks after treatment, animals show signs of disturbed glucose homeostasis but are not hyperglycemic, mimicking some aspects of the pre-symptomatic stage 2 of T1D(44). Our data show that at this stage islet microvessels, but not acinar vessels, present structural and phenotypical alterations of their cellular elements (pericytes and endothelial cells), resulting in functional defects that interfere with proper regulation of islet blood flow and vascular integrity. Our study thus supports islet microvascular dysfunction as an early event in diabetes pathogenesis in mice and humans(40).

We find in our study significant alterations in the density and phenotype of islet pericytes upon STZ treatment (**Figures 2,3**), particularly in the vicinity of inflamed endothelial cells (**Figure 2**). Islet pericytes have shown incredible morphological and phenotypic plasticity in rodent models of T1D(7, 35) and in models of hyperinsulinemia, insulin resistance and hypertension(26, 45–47). They hypertrophy and turn into ECM-producing myofibroblasts as shown by their close association with collagen fibrils (with some of these fibrils being present within their cytoplasm) and expression of myofibroblast markers such as periostin(26). A similar phenotypical switch of islet pericytes towards a myofibroblast-like cell is also observed upon multiple low-dose STZ treatment (**Figure 3**). Importantly, this phenotypical switch compromises pericyte function and vasomotor responses to vasoactive stimuli such as high glucose (**Figure 4**) and epinephrine (**Figure 5**). Abnormal vasomotor responses recorded *ex vivo* in living pancreas slices have repercussions *in vivo*, and compromise the acute increase in islet blood flow that occurs in response to glucose stimulation (functional hyperemia; **Figures 6,7**). Our data are in line with a previous report using contrast-enhanced ultrasound which revealed that islet blood flow dynamics was abnormal upon multiple low-dose STZ(20). In summary, STZ treatment modifies islet pericytes potentially interfering with their capacity to regulate local blood flow, particularly upon a glucose challenge.

Impaired islet hyperemic responses to glucose in this animal model could indicate that a functional beta cell mass is needed for this physiological response to occur. Beta cells in islets release ATP(48, 49), which can be degraded and converted into other purinergic signals (e.g. adenosine) by the action of different nucleotidases that are expressed by islet vascular and endocrine cells(50). Purinergic signals have vasoactive functions in different tissues such as the brain and heart(51, 52). Also in islets, we and others have shown that adenosine decreases islet pericyte [Ca^2+^]i and increases islet blood flow(13, 53). Besides adenosine, insulin itself also has important vasoactive actions throughout the body, acting not only on endothelial cells and triggering the release of different vasoactive molecules (e.g. nitric oxide, endothelin-1), but also directly affecting pericytes’ membrane potential and electrical activity(54). A decrease in functional beta cell mass upon STZ treatment would affect the levels of beta cell secretory products (e.g. insulin; ATP/adenosine) around islet vascular cells, impacting pericytes’ resting membrane potential and interfering with their electrical properties and capacity to control regional blood flow.

In this study, however, experiments were conducted two weeks after completing STZ treatment, when not only a decreased functional beta cell mass is present but also an inflammatory setting is well-established. A pro-inflammatory environment can contribute to myofibroblast appearance (a subset derived from pericytes; **Figure 3**) as these cells are highly responsive to cytokines and chemokines(37). Importantly, myofibroblasts (and fibroblasts) have different electrophysiological properties: unlike pericytes, these cells are not excitable and do not generate action potentials(55). An increase in their density could produce barriers to conduction of electrical signals through islet vascular beds, further compromising coordinated vascular responses that may be needed during acute changes of blood flow.

Pericyte dysfunction and their conversion into a myofibroblast-like cell may also involve their detachment from the capillary, interfering with pericytes’ vascular-stabilizing properties (56). Indeed, abundant literature exists supporting that a tight association of endothelial cells with pericytes is crucial for capillary homeostasis, as pericytes stabilize the microvasculature and control transendothelial permeability(10). Their conversion into myofibroblasts and detachment from the capillary wall can result in vulnerable vessels which are prone to instability and increased leakage, as we observe in islets from STZ-treated mice where high molecular weight (500kDa) dextran can leak from vessels in their vicinity (**Figure 7**). As it has been recognized in the field of pericyte biology, pericyte coverage of capillaries is not sufficient to ensure vascular stability, but “specific qualities of these cells are required”(10).

In summary, our study supports that in islets haemodynamic changes are present before the onset of diabetic symptoms. Determining their origin, examining their link with a defective beta cell mass and the contribution of a pro-inflammatory environment to these alterations will be needed for our understanding of the pathogenesis of diabetes. This knowledge would help designing strategies aimed at restoring the perivascular niche in islets, which we believe is crucial for proper endocrine cell function(11).

## Acknowledgements

The authors would like to thank Dr. Ruy Louzada (University of Miami) for careful revision of the manuscript, and Dr. Madina Makhmutova (University of Miami) for establishing the platform for imaging *in vivo* the exteriorized mouse pancreas.

## Author contributions

L.M.G performed immunohistochemistry, functional imaging with living pancreas slices, blood flow measurements, and analyzed data; I.S. analyzed blood flow data; E.P. performed immunohistochemistry; M.S.R adapted the Python analysis pipeline and analyzed blood flow data. J.A. designed the study, analyzed data and wrote the manuscript. All authors discussed the data, revised and contributed to the manuscript. Further information and requests for resources or reagents should be directed to the Lead Contact, Joana Almaça.

## Data Access and Responsibility

J.A. is the guarantor of this work and, as such, has full access to all the data included in the study and takes responsibility for the integrity of the data and the accuracy of the analysis. Any requests should be mailed to.

## Data and Resource Availability

All data generated or analyzed during this study are included in the published article (and its online supplementary files). The automated method to analyze blood flow is a resource made available in GitHub through this link: https://github.com/MSlakRupnik/Inst_Speed.

