## Supplementary material for "Microvascular homeostasis is compromised in pancreatic islets in a mouse model of beta cell loss and low-grade inflammation"

**Supplementary complete experimental procedures**

Multiple low-dose STZ mouse model

We administered the beta cell toxin streptozotocin (STZ; Sigma-Aldrich S0130) to mice as multiple consecutive doses of 50 mg/kg body weight for five days by intraperitoneal (i.p.) injections. STZ was dissolved in sodium citrate buffer (pH 4.5) right before injections. Equivalent volumes of sodium citrate buffer were injected in control animals (vehicle treated). All experiments were conducted from days 15-23 after treatment start (**Figure S1**). For *ex vivo* measurements of cytosolic Ca^2+^ levels in mural cells (pericytes and smooth muscle cells), we crossed mice that express Cre recombinase under the control of the mouse NG2 (*Cspg4*) promoter/enhancer (Jackson labs; strain #008533) with mice that express GCaMP3 downstream of a loxP-flanked STOP cassette (Jackson labs; strain #029043). Cre negative animals were used in all other experiments (*in vivo* imaging and immunohistochemistry). Adult mice (3-9 months old) from both genders were used. Glucose tolerance tests (2g/Kg BW; i.p.), following a six-hour fasting period, were performed 10 days after the last i.p. injection of STZ. Blood samples were collected from the tail and glucose measurements were made using a glucometer (Countour next EZ). All the experiments were conducted according to protocols and guidelines approved by the University of Miami Institutional Animal Care and Use Committee.

Preparation of living pancreas slices

Living pancreas slices were prepared from NG2-GCaMP3 mice as previously described (13). Briefly, mice were anesthetized with isoflurane followed by cervical dislocation. Low gelling temperature agarose (1.5%, Sigma Aldrich, cat. Nr. 39346-81-1, dissolved in HEPES-buffered solution without BSA) was injected in the common bile duct using a 30-gauge needle and 3 mL syringe. After injection, the tissue was cut in small pieces that were embedded in agarose and placed at 4°C for 10 min. Pieces of the pancreas were then sliced (150 μm thickness) in a vibrating blade microtome (VT1000S, Leica). To label blood vessels, a *Lycopersicon esculentum* derived fluorescent lectin (DyLight 649; Vector Labs, DL-1178-1) was injected (70 μL) in the tail vein 10 minutes before sacrifice.

Recording mural cell cytosolic Ca^2+^ responses and vasomotion in living pancreas slices

Living pancreas slices were placed on a coverslip in an imaging chamber (Wamer Instruments, Hamden, CT, USA) for imaging in an upright confocal microscope (upright Leica SP8 system). Slices were continuously perfused with HEPES-buffered solution containing 3 mM glucose and confocal images were acquired with LAS AF software (Leica Microsystems) using a 40X water immersion objective (NA 0.8). We used a resonance scanner for fast image acquisition to produce time-lapse volume recordings spanning 50-100 μm of the slice (z-steps: 4-8 μm; 512x512 XY pixels), at 5 s temporal resolution (XYZT imaging). GCaMP3 fluorescence was excited at 488 nm and emission detected at 510–550 nm, DyLight 649 labeled lectin was excited at 638 nm and emission detected at 650-680 nm. We recorded changes in the islet mural cell cytosolic Ca^2+^ levels ([Ca^2+^]i) and blood vessel diameter induced by glucose (16 mM) and epinephrine (10 μM – Sigma cat. nr.E4375) in islets (identified using the backscatter signal).

Using ImageJ sotware (<https://imagej.net/software/fiji/>), we examined changes in [Ca^2+^]i levels in mural cells (pericytes and SMCs) by drawing regions of interest around them and quantifying mean GCaMP3 fluorescence levels over time. Mean fluorescence intensity values were normalized to baseline levels in those same regions and plotted as ΔF/F (%). Islet pericyte [Ca^2+^]i responses are shown in the form of heatmaps generated using MatLab (<https://www.mathworks.com/products/matlab.html>). Pericytes were distinguished from SMCs by analyzing their morphology, shape of their cytoplasmic processes (elongated *versus* circular, respectively), location (within islet or covering larger vessels (diameter >10 μm) at the islet border or periphery) as previously published (38). Changes in vessel diameter were quantified as previously described (13, 38). For each stimulus, an average diameter value was calculated from ~10 diameter estimates obtained before stimulus application and at the end of the stimulus. To determine the extent of constriction/dilation, we pooled diameter data from different capillaries and islets for each group of mice and calculated the relative change in diameter (as % of baseline (3G) vessel diameter).

*In vivo* recording of blood flow and vascular leakage in the exteriorized pancreas

Animals (in fed state) were injected i.v. with 70 μL of fluorescent dextran (FITC-Dextran-500,000 Da – Sigma cat. nr. 46947 – at 20 mg/mL) before anesthesia. Animals were anesthetized with isoflurane and a small incision in their abdomen was made to exteriorize their pancreas. The exteriorized portion of the pancreas was placed bellow a 3D-printed vacuum window holding a glass coverslip (40) to keep the tissue steady under the confocal microscope (Nikon Upright AX R Confocal Imaging System; objective: CFI75 APO 25X (NA 1.1)). Islets were identified using the backscatter signal (excitation and emission at 638 nm) and FITC-dextran labeled vessels were visualized using a 488 nm laser and emission detected at 500-550 nm. We imaged islet blood vessels in single confocal planes at different times before (baseline videos) and after glucose (2 g or 4 g/Kg BW; i.p.) or epinephrine administration (0.9 mg/Kg BW; i.p.). Images were acquired at 15 frames/sec for 2 min every 5 minutes. Blood glucose levels were measured before and after isoflurane exposure and after glucose injection.

To assess the extent of dextran (500 kDa) extravazation from blood vessels towards the interstitial space in the islet vicinity, we quantified mean fluorescence intensities in the green channel in regions of interest (ROIs) drawn around the islet periphery excluding vessels, and divided values by fluorescence levels in islet blood vessels. For each islet, an average normalized dextran fluorescence in peri-islet interstititum was calculated.

Automated analysis of capillary blood flow using a Python pipeline

Videos of blood flowing through FITC-dextran labeled vessels in islets were captured at 15 frames per second. To quantify blood flow, we have adapted a set of interdependent tools to automatize as much as possible the analysis pipeline (41). Detailed pipeline analysis can be found on GitHub link: [https://github.com/MSlakRupnik/Inst_Speed](https://urldefense.com/v3/__https:/github.com/MSlakRupnik/Inst_Speed__;!!KVu0SnhVq1hAFvslES2Y!L5CCRZcWwnbjkzAdToZFgW8NWCouNWeToYWLsRSdE7dsf4go_TYhq9dI7qUD88nJemaZP6fgTthyfztVtwBwdyJCBjrj7_G_4xnN8Q$). In short, as we are mainly interested in capillary blood flow, for processing we chose the kernel size to approximately correspond to 5 µm, which is the average capillary diameter in islets. Detected events are fluorescent plasma, interrupted by dark periods caused by individual red blood cells (RBCs) moving through the blood vessel. An advantage of this approach is that it can automatically detect events occurring at any angle, even perpendicular to the optical plane, which significantly increases the detection rate in comparison to manual following in imageJ, which is accurate only in the plane parallel to the optical plane. To distill events (z-score > 3) from traces, we first performed a sequential filtering of traces at timescales starting from 0.5 s, and increasing by a factor as described previously, until the timescale of the longest event of interest was achieved. Each event has been characterized by a start time (t0), its maximal height, and the width at the half of the height (halfwidth, δt), which is our measurement of its duration (**Figure S5**). We detected an event at multiple timescales having similar starting time and comparable halfwidth (tolerance 20%). For a set of such events, a median halfwidth has been stored. The inverse value of the halfwidth of the single event within the ROI of a known size has been used to calculate the mean instant velocity (instantaneous speed; μm/s) of plasma passage through the corresponding capillary section, normalized to the frame rate of the video. The parameters we used to compare changes in blood flow were the instantaneous speed and the number of events collected for each ROI. We averaged these data for different capillaries for each islet and quantified absolute and relative changes in these parameters induced by glucose or epinephrine administrations for STZ- and vehicle-treated animals.

Immunohistochemistry

Two-three weeks after the first injection of STZ or vehicle (administered as 5 consecutive i.p. injections), mice were perfused with 4% PFA and their pancreata collected, embedded in sucrose (30% w/w), and frozen in Tissue-Tek Optimal Cutting Temperature (OCT) compound before cryosectioning (-20C). A cohort of vehicle- and STZ-treated mice were injected i.v. with 100 μL of lower molecular weight FITC-dextran (40,000 Da; lysine fixable; at 10mg/mL – Sigma cat nr. 2552718) and 30 min later perfused with PFA. Living pancreas slices used for physiological experiments ([Ca^2+^]i imaging) were immersed in 4% PFA for 1 hour and then washed with PBS. Pancreatic tissue sections (40 μm) or slices (150 μm) were incubated in blocking solution (PBS-triton X-100 0.3% and Universal Blocking Reagent; Biogenex, San Ramon, CA) for ~3 hours. Thereafter, sections were incubated for 48h (room temperature) with primary antibodies diluted in blocking solution. Slices and sections were immunostained for GFP (1:500, Abcam, cat. nr. Ab13970), pericyte marker neuron-glial antigen 2 (NG2, 1:100, Millipore, cat. nr. AB5320), alpha smooth muscle actin (αSMA; 1:250, Sigma, cat. nr. A5228), endothelial cell marker CD31 (1:25, Millipore, cat. nr. 550274), macrophage marker ionized Ca^2+^‐binding adaptor molecule 1 (Iba1; 1:250, Abcam, cat. nr. AB178846), adhesion molecule ICAM-1 (1:100, BD Pharmingen, cat. nr. 550287), the myofibroblast marker and ECM protein periostin (1:250, R&D, cat. nr. AF2955), Insulin (1:5, Dako, cat. nr. IR002) and Insulin-790 (1:250, Santa Cruz, cat. nr. Sc8033). Immunostaining was visualized by using Alexa Fluor conjugated secondary antibodies (1:500 in PBS; 16 h at 20 C; Invitrogen, Carlsbad, CA). Cell nuclei were stained with DAPI. Slides were mounted with Vectashield mounting medium (Vector Laboratories). Confocal images of immunostained sections or slides were acquired on an inverted White Light Laser confocal microscope (Leica Stellaris 5) with LAS AF software using a 63X oil immersion objective (NA 1.4) (Leica Microsystems).

Using ImageJ software, the area immunostained for different target antigens was quantified and compared to the total area of the region of interest: either inside the islet (islet density), around the islet (a ring of 20 μm was drawn around the islet to estimate peri-islet density) or within the surrounding acinar tissue (acinar density). Mander’s coefficients were determined to estimate the colocalization between pericyte markers (NG2 and PDGFRβ) and GFP or periostin, in confocal images of islets using the ImageJ plugin “Just Another Co-localization Plugin” (<https://imagej.nih.gov/ij/plugins/track/jacop2.html>).

Real-time PCR of sorted islet pericytes

Pancreatic islets were isolated from NG2-GCaMP3 treated with multiple low doses of STZ or with vehicle. After overnight culture, islets were digested for 5 min with trypsin (0.05 mg/mL) to obtain single cell suspensions. After few washes with cold PBS, dead cells were stained with DAPI (1:10000) and viable GFP-positive and GFP-negative cells were FACS sorted from cell suspensions in PBS + 1%FBS. Sorted cells were collected in lysis buffer and stored at -20°C. RNA from the different samples was extracted using RNAeasy micro kit (Quiagen), and stored at -80°C. RNA quality and purity was assessed using a Nanodrop (ND-1000 Spectrophotometer) and only RNA with a RNA integrity number (RIN) >7 was used further for reverse transcription. For cDNA synthesis, we used a high-capacity cDNA reverse transcription kit (Applied Biosystems) and 100 ng of RNA in each reaction (final volume 20 µL), following the manufacturer's protocols. Quantitative real-time PCR was performed using TaqMan fast universal PCR master mix and 10 ng of cDNA were used per reaction. We used TaqMan primers (FAM dye labeled) to determine expression of the following genes – *Rn18s, Cspg4, Acta2, PDGFRB, Cola1a, Postn, Ins1 and Gcg* - and run on a StepOnePlus real-time PCR system (Applied Biosystems). Data were analyzed using the ΔCT method. Briefly, expression of each gene was normalized to that of 18S ribosomal RNA (*Rn18s*) as a reference gene. To confirm that sorted GFP-positive cells were indeed pericytes, we compared the levels of pericyte genes (*Cspg4, Acta2, PDGFRb*) in this population with the levels of the same genes in GFP-negative cells (levels of *Ins* and *Gcg* were also examined in these samples), normalized to the internal reference *Rn18s* [using 2^(-ΔCT)], where ΔCT is the difference between the cycle threshold (CT) value of the gene of interest and the CT value of the reference *Rn18s*. Quantitative real-time PCR experiments were performed for 4 mice in each group.

Statistical analyses

For statistical comparisons we used Prism 10 (GraphPad software, La Jolla, CA) and performed unpaired t-tests when comparing data obtained for vehicle-treated with STZ-treated mice, or one-way ANOVA followed by Tukey’s multiple comparisons test when comparing different regions/conditions in vehicle- vs STZ. Multiple unpaired t-tests were applied when comparing data obtained at different time points in vehicle *versus* STZ-treated mice. One sample t tests were used to determine if average changes in vessel diameter induced by 16G or epinephrine were significantly different from 0. *p* values < 0.05 were considered statistically significant (indicated with an * in figures). Throughout the manuscript we present data as individual data points, mean ± SEM (bar graphs) or in box-and-whiskers plots where whiskers reflect minimum and maximum values.

**Supplementary Figure Legends**

**Supplementary Figure 1**

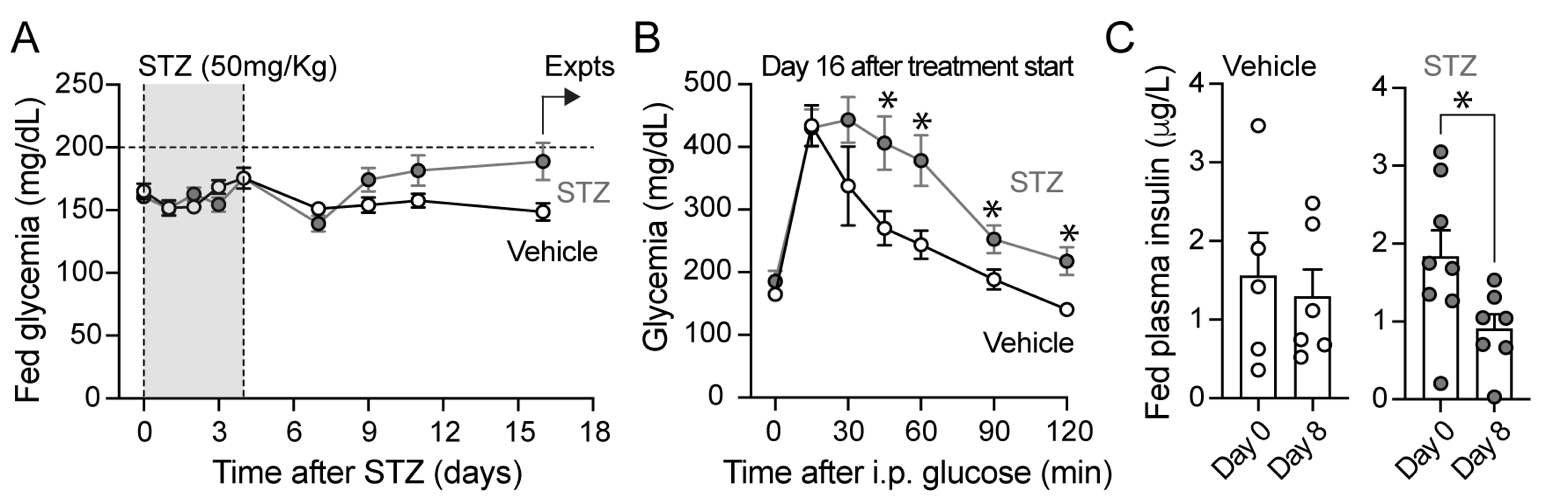

**Supplementary Figure 1. Glucose metabolism in mice treated with multiple low doses of STZ**

(A) Random fed glycemia measurements of all animals included in the study during and after vehicle- or STZ-treatment. STZ was administered i.p. as multiple consecutive doses of 50 mg/kg body weight for five consecutive days. Vehicle-treated animals received sodium citrate buffer. All experiments were conducted from days 15-23 after treatment start. (B) Glucose tolerance test (glucose dose of 2g/kg body weight; i.p.) conducted two weeks after STZ treatment had started (n=9-14 mice; multiple unpaired t test; **p*< 0.05). (C) Plasma insulin levels in the fed state collected before and 8 days after treatment start (paired t-tests; **p*< 0.05 for STZ-treated mice).

**Supplementary Figure 2**

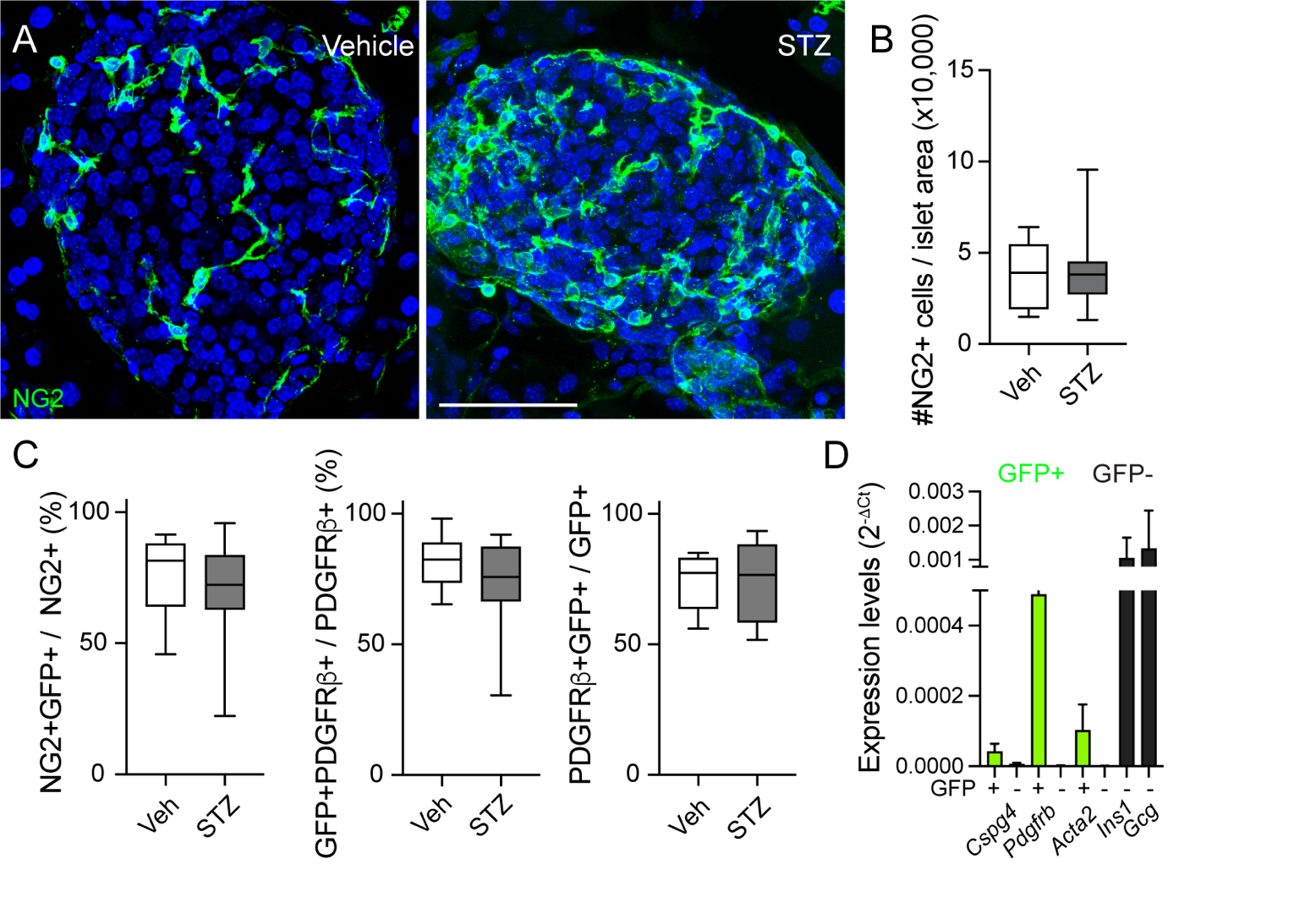

**Supplementary Figure 2. Pericyte number, phenotype and marker expression**

(A) Z-projections of confocal images of islets in pancreas sections from vehicle- and STZ-treated mice immunostained for pericytes (NG2; green). Scale bar = 40 μm. (B) Quantification of the number of NG2-positive cells in islets from vehicle- or STZ-treated mice, normalized to an average islet area of 10,000 μm^2^. (C) Mander coefficients estimating colocalization between GFP and pericyte markers NG2 and PDGFRβ in islets in pancreas sections from NG2-GCaMP3 mice. (D) Isolated islets from NG2-GCaMP3 mice were digested with trypsin to make single cell suspensions and subjected to FACS sorting. GFP-positive cells and GFP-negative cells were collected for RNA extraction. By real-time PCR, we examined the expression levels of genes encoding pericyte markers (*Cspg4*, *Pdgfrb* and *Acta2*) and islet hormones (*Ins1* and *Gcg*) in GFP-positive (+; green) and GFP-negative cells (-; black). mRNA levels were normalized to transcript levels of *Rn18s*.

**Supplementary Figure 3**

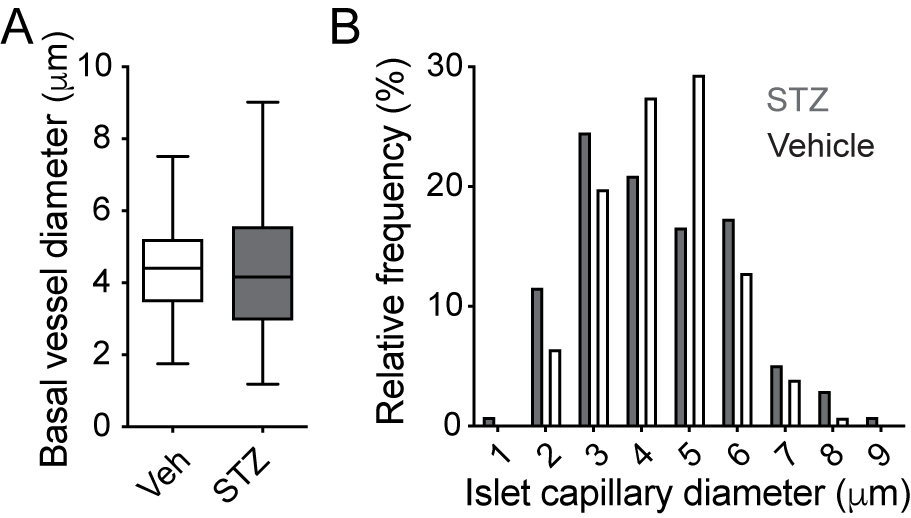

**Supplementary Figure 3. Basal islet capillary diameter in mice treated with multiple low doses of STZ**

(A) Quantification of diameters of islet capillaries (labeled with lectin) in mice treated either with vehicle- or STZ (unpaired t-test; *p*=0.4). Values measured under basal (3 mM) glucose concentration. (B) Histogram showing the percentage of vessels with different diameters in STZ- (gray bars) and vehicle-treated mice (white bars) under basal, non-stimulatory conditions.

**Supplementary Figure 4**

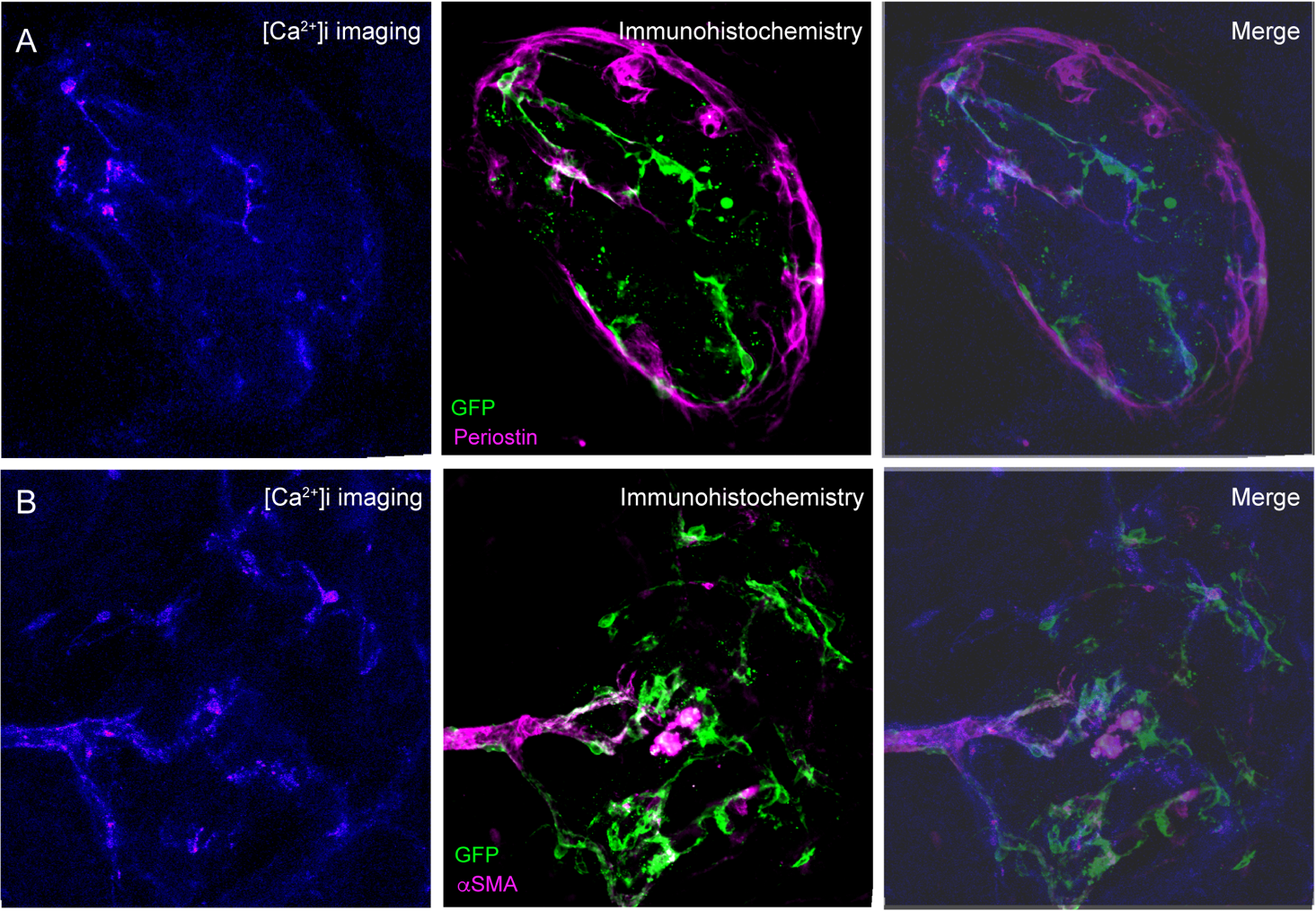

**Supplementary Figure 4. Correlating islet pericyte [Ca^2+^]i responses with their phenotype**

(A,B) Slices from NG2-GCaMP3 mice were used for physiology and subsequent immunostaining to match physiological responses of islet pericytes with their phenotype. Pericyte [Ca^2+^]i responses to 16G (A) or epinephrine (B) were recorded and the same slice was then fixed and immunostained for GFP (green) and periostin (magenta; A) or αSMA (magenta; B). Pericyte [Ca^2+^]i responses are shown in a pseudo color scale.

**Supplementary Figure 5**

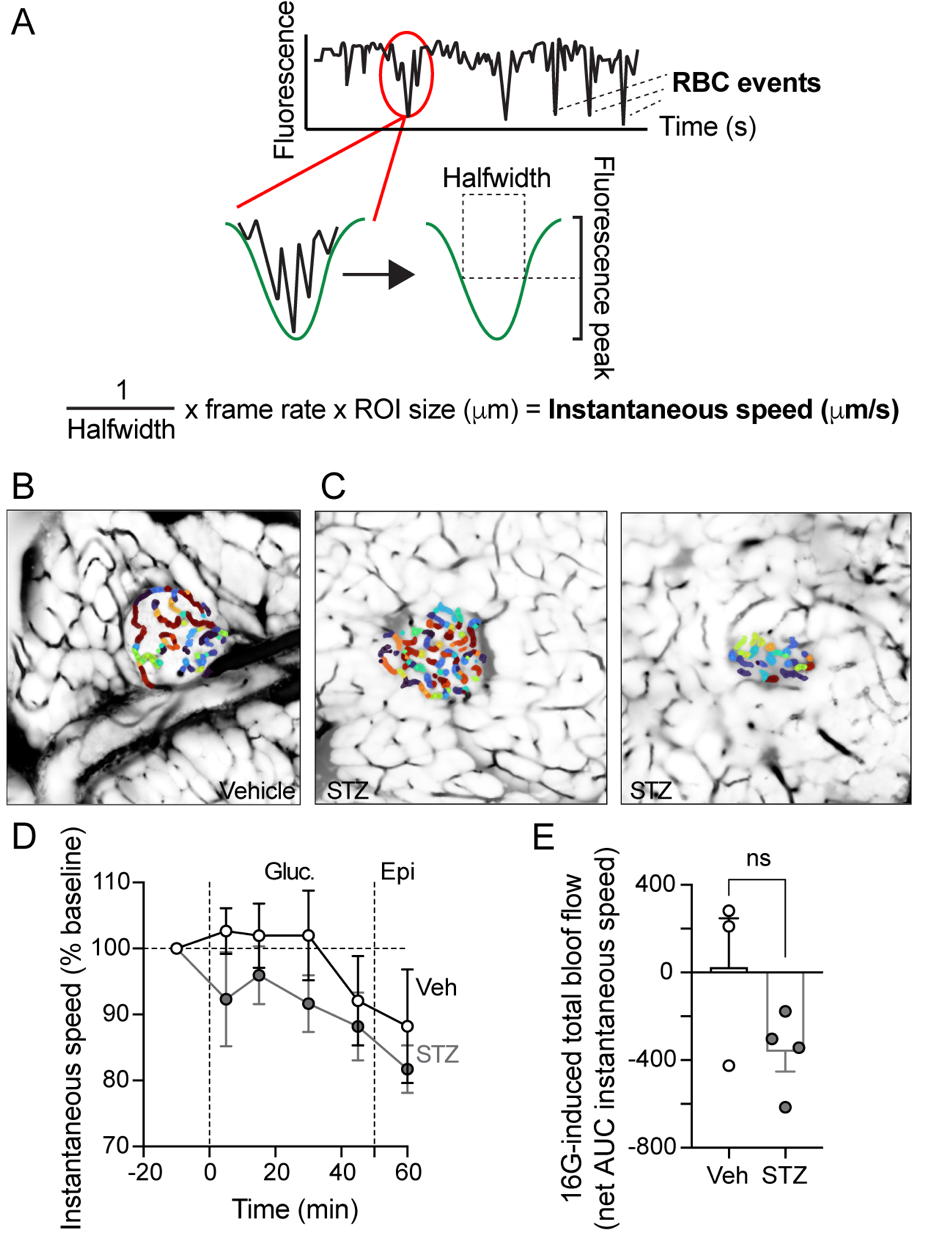

**Supplementary Figure 5. Quantification of blood flow in the exteriorized mouse pancreas using a Python-based pipeline**

(A) To quantify blood flow, we used a Python-based pipeline (GitHub link: [https://github.com/MSlakRupnik/Inst_Speed](https://urldefense.com/v3/__https:/github.com/MSlakRupnik/Inst_Speed__;!!KVu0SnhVq1hAFvslES2Y!L5CCRZcWwnbjkzAdToZFgW8NWCouNWeToYWLsRSdE7dsf4go_TYhq9dI7qUD88nJemaZP6fgTthyfztVtwBwdyJCBjrj7_G_4xnN8Q$)), which detects movement of individual red blood cells (RBCs) by quantifyfing over time changes in plasma fluorescence interrupted by dark periods caused by moving cells through the vessel (red oval). Each event is characterized by a start time (t0), its maximal height, and the width at the half of the height (halfwidth, δt), which is our measurement of its duration. The inverse value of the halfwidth of each single event within the ROI of a known size has been used to calculate the mean instantaneous speed (μm/s), normalized to the frame rate of the video (15frames/s). (B,C) Images showing automatically detected ROIs in capillaries in vehicle- (B) and STZ-treated islets (C). (D) Quantification of relative changes in instantaneous speed (shown as a % of baseline speed) of blood flowing through capillaries in acinar tissue surrounding islets in mice treated with STZ (n=4 mice; gray) or vehicle (Veh; n=3 mice; black). Values were measured at 5, 15, 30, and 45 minutes after intraperitoneal injection of 40% glucose (4 g/kg glucose i.p.), and 10 minutes post epinephrine (n=3-4 mice; multiple unpaired t test; *p*> 0.05). (E) Quantification of the net AUC of traces as those shown in (D) reflecting changes in instantaneous speed in capillaries in the acinar tissue (unpaired t-test; *p*>0.05).

**Supplementary movie legends**

**Supplementary Movie 1. Recording islet blood flow before dextrose injection in a control mouse**

Series of confocal images of islet blood vessels labeled with FITC-dextran (150 kDa; i.v.) in a control mouse pancreas exteriorized for *in vivo* imaging, before dextrose injection (gycemia after isoflurane anesthesia was 241 mg/dL). RBCs can be seen as shadows crossing vessel lumen. Images were acquired every 0.068 sec. Movie rate = 15 frames/s.

**Supplementary Movie 2. Recording islet blood flow 30 min after dextrose injection in a control mouse**

Series of confocal images of islet blood vessels labeled with FITC-dextran (150 kDa; i.v.) in a control mouse pancreas exteriorized for *in vivo* imaging, 30 min after dextrose (2 g/kg) i.p. injection (gycemia after glucose was 521 mg/dL). RBCs can be seen as shadows crossing vessel lumens at higher speed (comparison with Movie S1). Images were acquired every 0.068 sec. Movie rate = 15 frames/s.

**Supplementary Movie 3. Recording blood flow in the pancreas before dextrose injection in a mouse two weeks after STZ administration**

Series of confocal images of islet blood vessels labeled with FITC-dextran (500 kDa; i.v.) in the pancreas of a STZ-treated mouse. Images were acquired before dextrose injection (gycemia after isoflurane anesthesia was 307 mg/dL). Note movement of RBCs through vessels in endocrine and exocrine tissue, as well as leakage of dextran towards tissue interstitial space. Movie rate = 15 frames/s.

**Supplementary Movie 4. Recording blood flow in the pancreas after epinephrine in a mouse two weeks after STZ administration**

Series of confocal images of islet blood vessels labeled with FITC-dextran (500 kDa; i.v.) in the pancreas of a STZ-treated mouse. Images were acquired after epinephrine injection (0.9 mg/Kg BW; i.p.). Movement of RBCs through vessels in endocrine and exocrine tissue decreases, while leakage of dextran towards tissue interstitial space increases (comparison with Movie S3). Movie rate = 15 frames/s.
